# The dynamic three-dimensional organization of the diploid yeast genome

**DOI:** 10.1101/091827

**Authors:** Seungsoo Kim, Ivan Liachko, Donna G Brickner, Kate Cook, William S Noble, Jason H Brickner, Jay Shendure, Maitreya J Dunham

**Affiliations:** Department of Genome Sciences, University of Washington, Seattle, WA; Department of Molecular Biosciences, Northwestern University, Evanston, IL; Howard Hughes Medical Institute, University of Washington, Seattle, WA

## Abstract

The budding yeast *Saccharomyces cerevisiae* is a long-standing model for the three-dimensional organization of eukaryotic genomes. Even in this well-studied model, it is unclear how homolog pairing in diploids and environment-induced gene relocalization influence overall genome organization. Here, we performed high-throughput chromosome conformation capture on diverged *Saccharomyces* hybrid diploids to obtain the first global view of chromosome conformation in diploid yeasts. After controlling for the Rabl-like orientation, we observe significant homolog proximity that increased in saturated culture conditions. Surprisingly, we observe a localized increase in homologous interactions between the *HAS1* alleles specifically under galactose induction and saturated growth, mediated by association with nuclear pore complexes at the nuclear periphery. Together, these results reveal that the diploid yeast genome has a dynamic and complex 3D organization.

## Introduction

The genome is actively organized in the nucleus, in both space and time, and this organization impacts fundamental biological processes like transcription, DNA repair, and recombination (Taddei et al., 2010). The budding yeast *S. cerevisiae* has been a useful model for studying eukaryotic genome conformation and its functional implications (Taddei et al., 2010). The predominant feature of yeast 3D genome organization is its Rabl-like orientation (Jin et al., 1998) (Figure 1A): during interphase, the centromeres cluster at one end of the nucleus, attached to the spindle pole body, and chromosome arms extend outward toward the nuclear periphery where the telomeres associate (Therizols et al., 2006), like in anaphase. In addition, the ribosomal DNA array forms the nucleolus, opposite the spindle pole (Yang et al., 1989), splitting chromosome XII into two separate domains that behave as if they were separate chromosomes. This organization largely persists through the cell cycle (Jin et al., 1998) and even in stationary phase, albeit with increased telomere clustering and decreased centromere clustering (Guidi et al., 2015; Rutledge et al., 2015).

**Figure 1.**
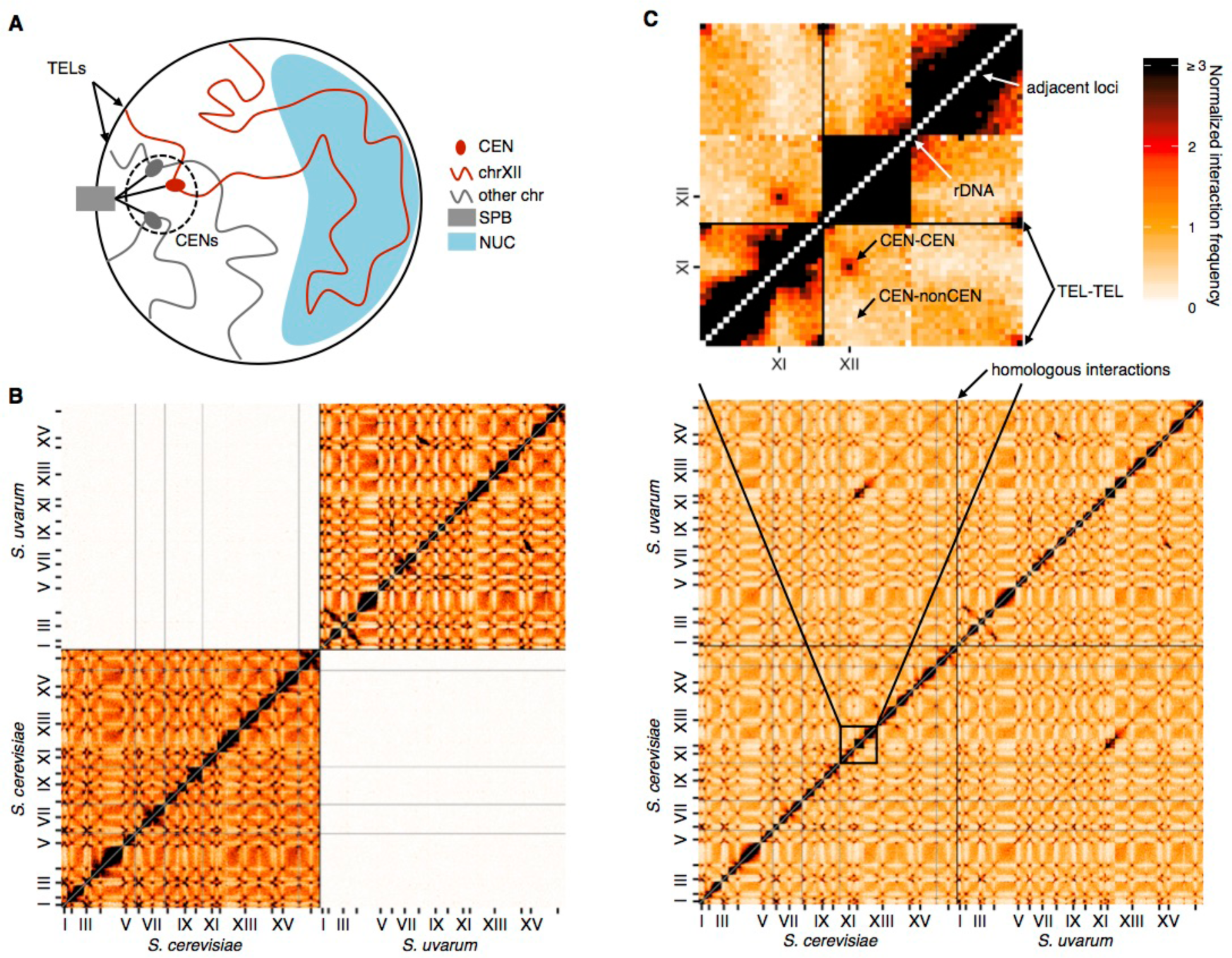
Diverged hybrids provide a genome-wide view of diploid chromosome conformation. (**A**) Schematic of the Rabl-like orientation. CEN, centromere; SPB, spindle pole body; TEL, telomere; NUC, nucleolus. (**B**) Hi-C contact map for saturated *S. cerevisiae* and *S. uvarum* mixture control, at 32 kb resolution. Each axis represents the *S. cerevisiae* genome followed by the *S. uvarum* genome in syntenic order, separated by a black line. Ticks indicate centromeres. Odd-numbered centromeres are labeled. Rows and columns with insufficient data are colored grey. (**C**) Hi-C contact map for saturated *S. cerevisiae* x *S. uvarum* hybrid, as in (**B**). A portion of the map, outlined in black, is enlarged above with annotated features of the Rabl-like orientation.

Genome-wide chromosome conformation capture methods like Hi-C have both confirmed these microscopy observations and permitted systematic analyses of the functional clustering of genomic elements like tRNA genes and origins of replication (Duan et al., 2010). However, multiple studies have argued that a simple volume exclusion polymer model of chromosomes in a Rabl-like orientation is sufficient to explain microscopy and Hi-C data of the budding yeast genome (Tjong et al., 2012; Wong et al., 2012), at least in haploids grown under standard lab conditions. Even the clustering that is observed may simply be a consequence of bias in chromosomal positions rather than active molecular interactions (Rutledge et al., 2015; Tjong et al., 2012).

Although this simplicity is attractive, diploidy and variable environmental conditions may add complexity to yeast genome conformation. In diploid yeast, homologous chromosomes have been observed to pair during mitotic growth (Burgess and Kleckner, 1999; Burgess et al., 1999; Dekker et al., 2002), as they do in *Drosophila* (Metz, 1916), but the extent of this pairing has been debated (Lorenz et al., 2003). In both haploid and diploid yeast, genome conformation can also change in response to environmental conditions, but systematic studies of such changes have been lacking.

## Results

We performed Hi-C on interspecific hybrids between diverged *Saccharomyces* species to obtain the first genome-wide view of chromosome conformation in diploid yeasts. The high sequence similarity of homologous chromosomes in diploid *S. cerevisiae* precludes observation of interactions between them using sequencing-based methods. However, divergent *Saccharomyces* species can form stable hybrids (González et al., 2006; Mertens et al., 2015), e.g. between *S. cerevisiae* and *S. paradoxus* (90% nucleotide identity in coding regions (Kellis et al., 2003)) or its more distant relative *S. uvarum* (also known as *S. bayanus* var. *uvarum*; 80% nucleotide identity in coding regions (Kellis et al., 2003)). These interspecific hybrids are sufficiently diverged to allow straightforward sequence-level discrimination of homologs of homologs (**Figure 1—figure supplement 1A**) but have maintained nearly complete synteny (Fischer et al., 2000). For comparison, we also analyzed hybrids between *S. cerevisiae* strains Y12 and DBVPG6044, which are much less diverged (~99% nucleotide identity) (Liti et al., 2009). We confirmed the minimal impact of mapping and experimental artifacts by mapping Hi-C data from each individual species or strain (**Figure 1—figure supplement 1B-D**) and mixtures thereof (**Figure 1B, Figure 1—figure supplement 2**) to the hybrid reference genomes.

The most prominent features of Hi-C data from diploid yeast are the signatures of a Rabl-like orientation (Figure 1C). As in all Hi-C datasets, the contact map exhibits a strong diagonal signal indicating frequent intrachromosomal interactions between adjacent loci. In addition, pericentromeric regions interact frequently with one another, but infrequently with regions far from centromeres, as expected from the clustering of centromeres at the spindle pole body. Telomeric regions also preferentially interact, consistent with their clustering at the nuclear periphery. Finally, the rDNA-carrying chromosomes each behave as two separate chromosomes divided by the nucleolus, with frequent interactions on either side of the rDNA array but not across it.

In addition to these previously described phenomena, we observed an off-diagonal line of increased interaction suggestive of homolog pairing (Figure 1C). In microscopy (Burgess et al., 1999), recombination efficiency (Burgess and Kleckner, 1999), and chromatin conformation capture (Dekker et al., 2002) assays, homologous loci tend to be closer together than nonhomologous loci. This has been suggested to be the result of transient pairing between homologous nucleosome-free DNA, which may function to prepare the genome for meiosis or homology-directed repair. However, it has also been suggested that the observation of homolog proximity is an artifact of the Rabl-like orientation or FISH methods (Lorenz et al., 2003). This debate remains unresolved in part due to the targeted nature of previous studies, wherein each pair of homologous loci is only compared to a limited number of nonhomologous loci.

To systematically investigate whether homolog proximity can be explained by the Rabl-like orientation, we compared our experimental data from *S. cerevisiae* x *S. uvarum* hybrids to simulated data from a volume exclusion polymer model of the Rabl-like orientation. This model did not explicitly encode homolog pairing (Tjong et al., 2012) and served as a negative control to calibrate our quantification. We quantified homolog proximity by comparing the frequency of each interaction between a pair of homologous loci to the set of nonhomologous interactions involving either locus (**Figure 2—figure supplement 1**). Initially, we controlled for the Rabl-like orientation by restricting comparisons to interactions with loci at a similar distance from the centromere (at a resolution of 32 kb), as previous studies have done (Burgess et al., 1999; Lorenz et al., 2003). Using this approach, we find that the polymer simulation predicts strong homolog proximity (**Figure 2A**). Thus, the long-used approach of comparing homologous interactions to nonhomologous interactions at the same centromeric distance may not fully account for the Rabl-like orientation. Polymer models suggest that short chromosomes interact preferentially, due to their dual telomeric tethering at the nuclear periphery and centromeric tethering at the spindle pole (Tjong et al., 2012); therefore, we further restricted comparisons to loci on chromosome arms of similar length (within 25%). This additional restriction dramatically reduced the signal of homolog proximity for the polymer model, but not the experimental data (**Figure 2B**).

**Figure 2.**
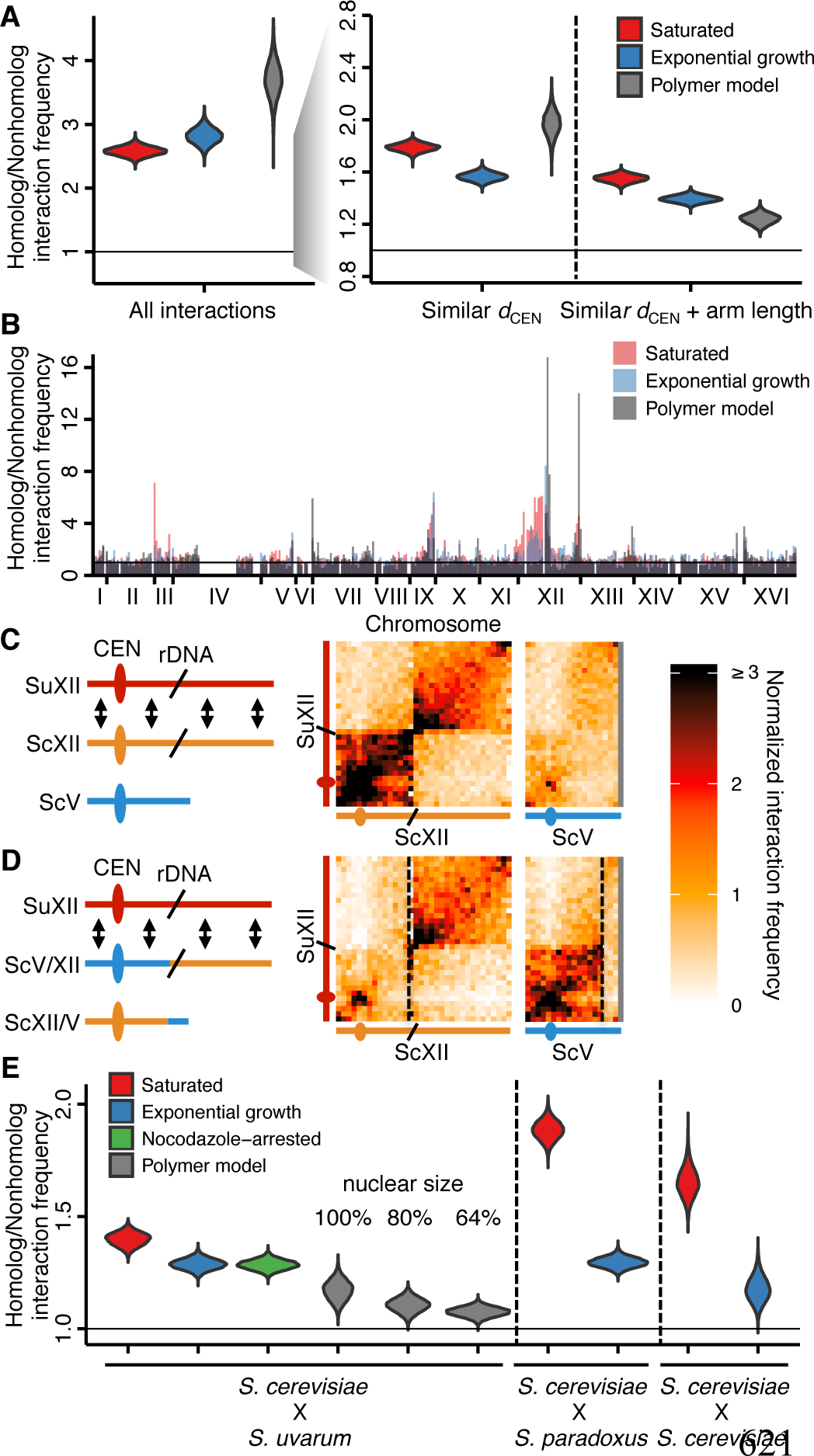
Homolog proximity exceeds predicted effects of Rabl-like orientation. (**A**) Estimates of homolog proximity (ratio of homologous to nonhomologous interaction frequencies) throughout the *S. cerevisiae* x *S. uvarum* hybrid genome as a function of increasing comparison stringency (left to right) to account for Rabl-like orientation (**Figure 2—figure supplement 1**). Saturated culture data are shown in red, exponential growth in blue, and simulated data from a homology-agnostic polymer model in grey, *d*_*CEN*,_ distance from centromere. (**B**) Variation in homolog proximity across the *S. cerevisiae* x *S. uvarum* hybrid genome at 32 kb resolution, in saturated culture (red), exponential growth (blue), and the polymer model (grey). Nonhomologous interactions were restricted to similar centromere distance and chromosome arm length. Data are plotted by *S. cerevisiae* genome position, x ticks indicate ends of chromosomes. (**C** and **D**) Schematics and Hi-C contact maps (at 32 kb resolution) of interactions between *S. uvarum* chromosome XII (SuXII) and either *S. cerevisiae* chromosome XII (ScXII) or *S. cerevisiae* chromosome V (ScV), in wild-type *S. cerevisiae* x *S. uvarum* hybrids (**C**) and a strain with a translocation between ScXII and ScV (**D**), both in saturated cultures. Ovals indicate centromeres and slanted lines indicate rDNA arrays. Double-headed arrows indicate enhanced interactions. Dashed lines indicate translocation breakpoints. (**E**) Estimates of homolog proximity across conditions, polymer models, and hybrids. Calculated as in (**A**), but excluding chromosome XII and all centromeres. Saturated culture data are shown in red, exponential growth in blue, nocodazole-arrested in green, and polymer models in grey.

Comparing homolog proximity across the genome, we noticed extensive interactions between the homologous chromosomes carrying the rDNA arrays (**Figure 2B**). To test whether this enrichment for interactions is due to sequence-dependent homolog pairing, we created a translocation that swapped most of the centromeric half of *S. cerevisiae* chromosome XII with an equivalently sized portion of *S. cerevisiae* chromosome V. In this translocated strain, interactions between *S. uvarum* chromosome XII and *S. cerevisiae* chromosome V are enriched instead of *S. cerevisiae* chromosome XII (**Figure 2C,D and Figure 2—figure supplement 2A,B**), suggesting that homolog proximity of chromosomes carrying the rDNA arrays is due to the presence of the rDNA rather than the particular sequence of the chromosome that carries it. We propose that the rDNA-carrying chromosomes are uniquely positioned within the nucleus due to their tethering at the nucleolus(Duan et al., 2010) (**Figure 2—figure supplement 2C**). This shared tethering would then cause enhanced interactions between the homologous proximal and distal segments of these chromosomes and inflate the signal for apparent homology-dependent pairing.

Based on these findings, we excluded the rDNA-carrying chromosomes from estimates of homolog proximity. Even with these stringent constraints, we find that the observed interaction between homologous alleles exceeds that predicted simply due to the Rabl-like orientation (**Figure 2E**). The left arm of chromosome III and the right arm of chromosome IX exhibit particularly strong homolog proximity (**Figure 2B**).

In all hybrids, homolog proximity is substantially greater in saturated cultures approaching stationary phase than in exponential growth (**Figure 2E**), consistent with previous observations (Burgess et al., 1999). One explanation for this result is a difference in the strength of sequence-dependent homolog pairing. However, this difference could also be a consequence of the reduced cell cycling coupled with loss of homolog proximity during G2-phase (Burgess et al., 1999) or smaller nuclear size in cells approaching stationary phase (Guidi et al., 2015). To test the first hypothesis, we performed Hi-C on nocodazole-arrested (G2-phase) cells, which were previously reported to exhibit reduced homolog proximity (Burgess et al., 1999). If increased homolog proximity in saturated cultures were due to a lack of cells with decreased homolog proximity in G2-phase, arrested cells should exhibit even less homolog proximity than exponentially growing cells. However, we find that nocodazole arrest does not substantially reduce homolog proximity (**Figure 2E**).

We next sought to evaluate the effect of nuclear size on homolog proximity. We compared alternate versions of our polymer model of the Rabl-like orientation with proportionally smaller nuclei, at 80% and 64% of the original size. In these models, smaller nuclear size led to decreased homolog proximity (**Figure 2E**). This approach cannot assess the effect of nuclear size on sequence-dependent homolog pairing, which is absent from our models. Nevertheless, these models suggest that the difference in homolog proximity between saturated and exponentially growing cultures cannot be explained by the effect of nuclear size on homolog proximity driven by the Rabl-like orientation, and provide additional support for homolog pairing beyond the Rabl-like orientation.

We also searched our dataset for evidence of highly specific changes in genome conformation at the scale of individual genes. Microscopy studies have revealed inducible genes that relocate to the nuclear periphery upon activation due to association with nuclear pores (e.g. *GAL1* (Brickner et al., 2016; Casolari et al., 2004; Dultz et al., 2016), *INO1* (Brickner and Walter, 2004), *HXK1* (Taddei et al., 2006), *TSA2* (Ahmed et al., 2010), *HSP104* (Dieppois et al., 2006)), which can increase (Ahmed et al., 2010; Brickner and Walter, 2004; Brickner et al., 2016; Taddei et al., 2006) gene expression. Although DNA interactions with components of the nuclear pore complex (NPC) have been identified genome-wide by chromatin immunoprecipitation (Casolari et al., 2004), it remains unclear whether relocalization of specific genes impacts global genome conformation.

We first focused on the galactose metabolism gene *GAL1.* This gene and its neighbors *GAL7* and *GAL10* move upon galactose induction from their location near the spindle pole body to a nuclear pore complex at the nuclear periphery (Casolari et al., 2004; Dultz et al., 2016) (**Figure 3A**). Consistent with this expectation, we found using Hi-C that both *GAL1* loci interacted less with pericentromeric regions upon galactose induction (**Figure 3B-D**). Despite previous reports that the homologous *GAL1* loci preferentially interact with each other during galactose induction (Brickner et al., 2016), we do not see a clear signal for increased pairing (**Figure 3—figure supplement 1**), perhaps because of the high basal interaction frequency between pericentromeric loci.

**Figure 3.**
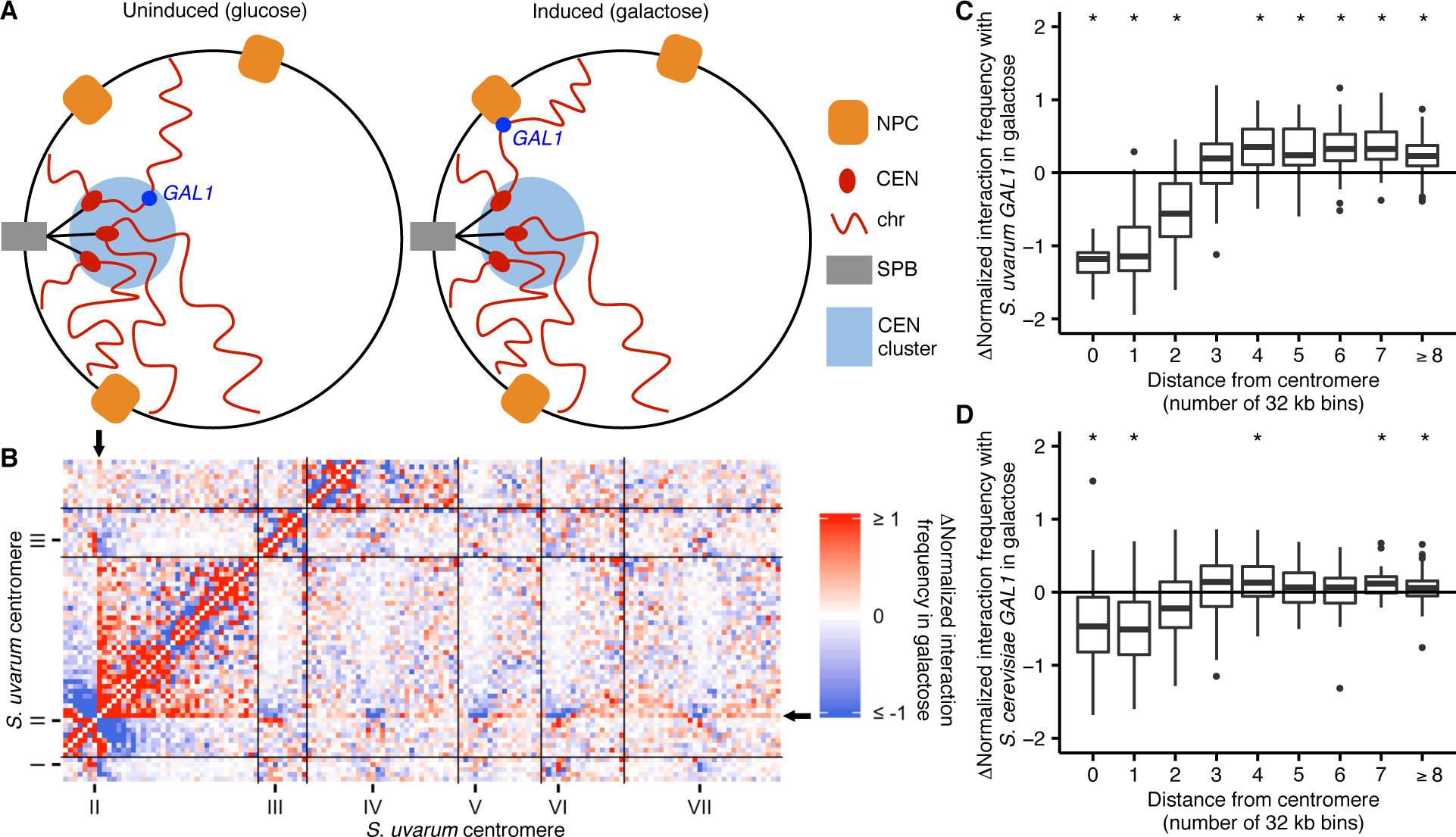
*GAL1* shifts away from centromeres upon galactose induction. (**A**) Schematic of *GAL1* positioning (dark blue) in glucose (left) and galactose (right). NPC, nuclear pore complex; CEN, centromere; chr, chromosome; SPB, spindle pole body. (**B**) Example region of differential Hi-C map of *S. cerevisiae* x *S. uvarum* hybrids in galactose vs. glucose, at 32 kb resolution. Interactions that strengthen in galactose are in red, while those that weaken are in blue. Ticks indicate centromeres; black lines indicate chromosomes. Arrows indicate location of *S. uvarum GALL* (**C**) Boxplot of the difference in *S. uvarum GAL1* interaction frequency in galactose vs. glucose across the *S. cerevisiae* x *S. uvarum* genome, excluding intrachromosomal interactions and binned by distance from the centromere (in 32 kb bins). Whiskers correspond to the highest and lowest points within the 1.5 x interquartile range. *\*P <* 0.05 after Bonferroni correction (n = 9); Mann-Whitney test. Note: some outliers are beyond the plot range and are not shown. (**D**) Same as (**C**) for *S. cerevisiae GALL*

Having established that we could detect the known inducible relocalization of the *GAL1* gene, we looked for other specific changes in genome conformation in the well-studied environmental conditions of galactose induction and growth saturation (approaching stationary phase). Surprisingly, we observed dramatically increased interactions between homologous loci near the gene *HAS1* on chromosome XIII under both growth saturation and galactose induction, compared to standard exponential growth in glucose (**Figure 4A**). In fact, under inducing conditions this interaction is among the strongest genome-wide, excluding pericentromeric and subtelomeric regions (top interaction among over 83,000; Figure 4—figure supplement 1A,B). No known galactose-induced genes are in or near this region. Nevertheless, this inducible homolog proximity appears to be evolutionarily conserved, as it occurs in all three tested interspecific hybrids (**Figure 4A-C**).

**Figure 4.**
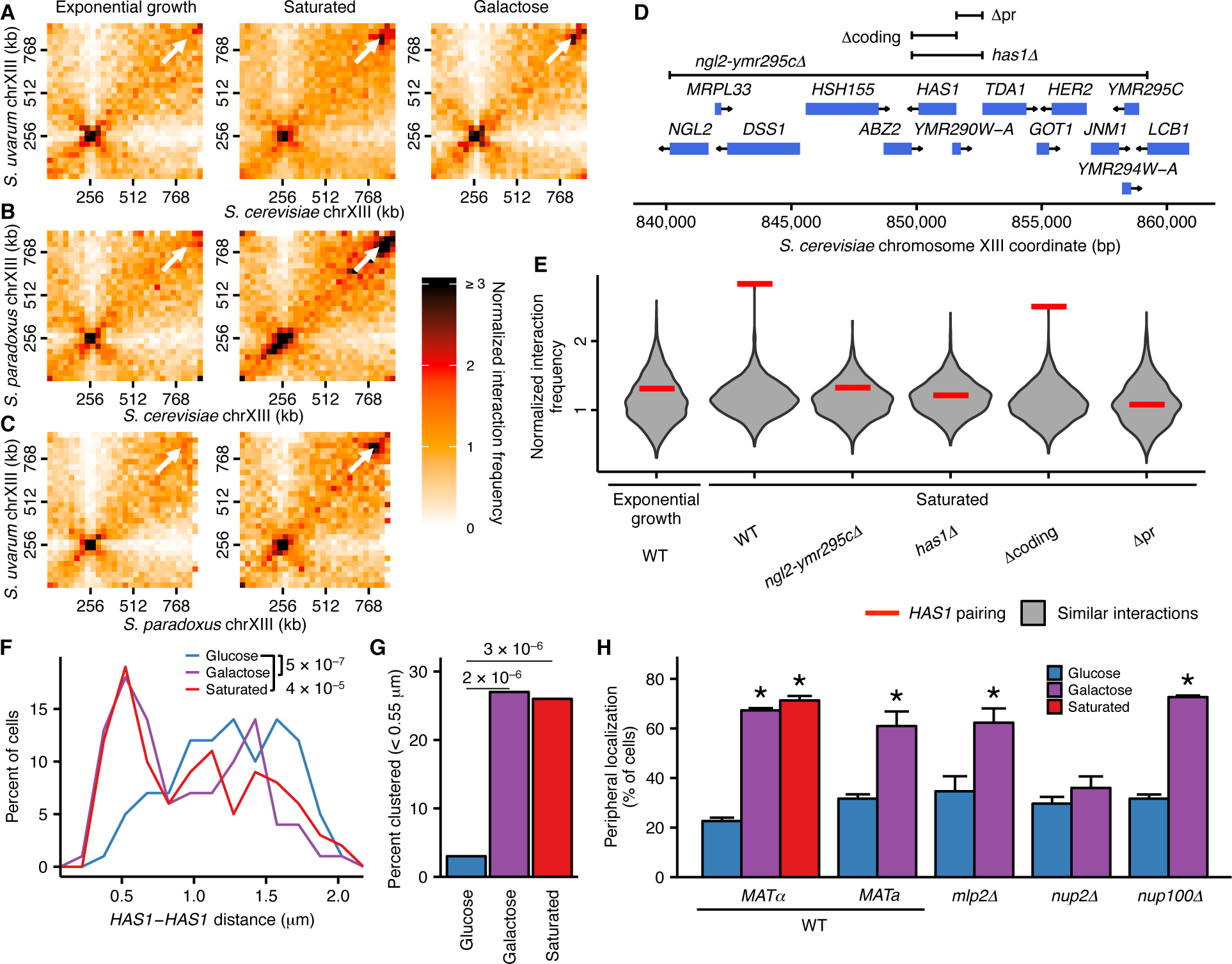
Inducible relocalization and pairing of *HAS1* homologs is evolutionarily conserved, sequence-specific, and mediated by nuclear pores. (**A, B**, and **C**) Hi-C contact maps of chromosome XIII interactions at 32 kb resolution in *S. cerevisiae* x *S. uvarum* (**A**), *S. cerevisiae* x *S. paradoxus* (**B**), and *S. paradoxus* x *S. uvarum* (**C**) hybrids in exponential growth (left column), saturated cultures (middle column), and in *S. cerevisiae* x *S. uvarum* hybrids (**A**), galactose (right column). White arrows indicate the interaction between the homologous *HAS1* loci. (**D**) Genome browser shot of open reading frames (ORFs; blue boxes) and tested deletions (brackets) in the *S. cerevisiae* region surrounding the gene *HAS1*, from positions 840,000–860,000 (**Figure 4—figure supplement 1A**). Arrows indicate ends and directionalities of ORFs. (**E**) Strength of *HAS1* pairing (red lines) compared to similar interactions (grey violin plots; i.e. interactions between an *S. cerevisiae* locus and an *S. uvarum* locus, where both loci are ≥ 15 bins from a centromere and ≥ 1 bin from a telomere, and not both on chromosome XII) in wild-type and deletion strains of *S. cerevisiae* x *S. uvarum.* (**F**) Distributions of the distance between the *HAS1* homologs measured by microscopy in *S. cerevisiae* diploids in glucose (blue), galactose (purple), and saturated cultures (red). *n* = 100 for each condition. P-values were calculated using the Wilcoxon rank sum test. (**G**) Frequency of *HAS1* alleles less than 0.55 μm apart, measured as in (**F**), in glucose (blue), galactose (purple), or saturated cultures (red). P-values were calculated using Fisher’s exact test. (**H**) Proportions of cells exhibiting peripheral *HAS1* localization in haploid *S. cerevisiae* in strains with and without deletions of nuclear pore components. Experiments were performed in biological triplicate, with *n* ≥ 30 per experiment. *P < .05, Student’s *t* test. Center values and error bars represent mean ± s.e.m.

To explore whether this pairing depends on the presence of specific sequences, we created various deletions of the *S. cerevisiae* copy of the region, ranging from a 20 kb region from *NGL2* through *YMR295C* (**Figure 4—figure supplement 1A**) to a single 1 kb intergenic region containing the promoters for *HAS1* and *TDA1 (HAS1pr-TDA1pr;* **Figure 4D**). Every deletion that included the *HAS1* promoter reduced the saturated *HAS1* homolog interaction frequency back to uninduced levels, indicating that this inducible pairing is in fact sequence-dependent (**Figure 4E**). In contrast, deletion of only the *HAS1* coding sequence had minimal impact, which shows that the deletion construct itself did not impede inducible pairing (**Figure 4E**). To test whether the *HAS1pr-TDA1pr* region is sufficient to produce inducible pairing, we moved the *S. cerevisiae* copy of this region to the left arm of *S. cerevisiae* chromosome XIV. Interestingly, the ectopic *HAS1pr-TDA1pr* allele exhibited inducible interactions with the *S. uvarum HAS1pr-TDA1pr*, though not to the same extent as the endogenous allele (**Figure 4—figure supplement 1C-E**). This further confirms the role of the *HAS1pr-TDA1pr* region in *HAS1* pairing, but suggests that other regions or possibly its chromosomal position may also contribute. To verify whether this pairing occurs in homozygous *S. cerevisiae* diploids in addition to diverged hybrids, we labeled both *HAS1* loci with integrated LacO arrays targeted by LacI-GFP and measured the distance between them under glucose, galactose, and saturation with microscopy. Consistent with our Hi-C data, the *HAS1* homologs were closer together in galactose-induced and saturated cultures than in glucose (**Figure 4F,G**).

Based on previous studies of relocalized genes (Brickner et al., 2012, 2016), we hypothesized that *HAS1* pairing might be mediated by interactions of both alleles with the nuclear pore complex (**Figure 4—figure supplement 1F**). Therefore, we tested whether the *HAS1* loci are relocalized to the nuclear periphery during galactose induction, using microscopy in haploid *S. cerevisiae.* Indeed, the *HAS1* locus shifted to the nuclear periphery upon galactose induction and in saturated culture conditions (**Figure 4H**). Prior genome-wide analysis of nuclear pore interactions in glucose and galactose (Casolari et al., 2004) had not identified the relocalization of the *HAS1* locus, likely due to limited coverage of noncoding sequences. To confirm whether this inducible reorganization was dependent on association with nuclear pores, we repeated our analysis in strains with deletions of nuclear pore components *NUP2* or *NUP100*, or pore-associated protein *MLP2.* As in other cases of gene relocalization, Nup2 but not Nup100 was required for peripheral localization of *HAS1.* However, unlike other relocalized genes (Ahmed et al., 2010; Brickner et al., 2016; Luthra et al., 2007), *HAS1* relocalization did not require Mlp2, suggesting a distinct mechanism of *HAS1* relocalization. Together, these data are consistent with inducible, sequence-specific pairing at the homologous *HAS1* loci mediated by transcription factor binding both within, and potentially also beyond, the *HAS1pr-TDA1pr* region upon induction. The *HAS1pr-TDA1pr-bound* transcription factor(s) could directly or indirectly associate with nuclear pores, leading to increased interaction between the homologous *HAS1* loci (**Figure 4—figure supplement 1G**).

## Discussion

The precise functional implications of genome conformation remain an open question, not only in budding yeast, but in all organisms. Although the budding yeast *S. cerevisiae* is thought to have a simple genome organization, it continues to serve as a useful model for phenomena like homolog interactions. Furthermore, our identification of a novel inducible conformational change in two highly studied conditions—a phenomenon that is conserved across highly diverged *Saccharomyces* species—suggests that we still have more to learn even about yeast genome organization. Admittedly, Hi-C has limited resolution (32 kb in our study) and may not be able to detect some types of gene relocalization. Nevertheless, Hi-C analysis of diverged hybrids allowed the first genomic analysis of diploid genome conformation and may add information orthogonal to microscopy and chromatin immunoprecipitation. Together, Hi-C, microscopy, and genetic perturbations will allow us to advance our understanding of genome conformation and its function.

## Materials and Methods

### Strain construction

All yeast strains used in this study are listed in **Supplementary File 1**.

Hybrid strains were created by mating haploid strains and auxotrophic selection according to standard protocols.

The ScV-ScXII translocation *S. cerevisiae* x *S. uvarum* strain was generated by first creating the translocation in the haploid *S. cerevisiae* strain BY4742, followed by mating with *S. uvarum.* A cassette containing *hphMX* followed by the first half of *URA3*, an artificial intron, and a *lox71* site was amplified from pBAR3 (Levy et al., 2015) and integrated into the intergenic region between *YLR150W* and *YLR151C.* A second cassette containing a *lox66* site, an artificial intron, the second half of *URA3*, and *natMX* was amplified from pBAR2-natMX (pBAR2 (Levy et al., 2015) with *natMX* in place of *kanMX)* and integrated into the intergenic region between *YER151C* and *YER152C.* The translocation was induced by transforming the resulting strain with the galactose-inducible Cre plasmid pSH47-kanMX (pSH47 (Güldener et al., 1996) with *kanMX* in place of *URA3)*, and then inducing Cre recombination by plating on YP + galactose medium. Successful translocation strains were selected by growing in medium lacking uracil, and verified by PCR across the translocation junctions. This *S. cerevisiae* strain was then mated with *S. uvarum* strain ILY376.

Deletion strains were made in *S. cerevisiae* x *S. uvarum* hybrids, by replacing regions of interest with the *hphMX* cassette. Strains were verified by PCR across each deletion junction.

The knock-in strain was made by integrating the *HAS1pr-TDA1pr* region followed by the *natMX* cassette into the region between *YNL266W* and *YNL267W (PIK1)* on *S. cerevisiae* chromosome XIV in the *S. cerevisiae* x *S. uvarum* hybrid YMD3269 *(HAS1pr-TDA1pr* deletion).

Plasmids pAFS144 (Straight et al., 1996), p5LacIGFP (Randise-Hinchliff et al., 2016), pER04 (Randise-Hinchliff et al., 2016) and pFA6a-kanMX6 (Longtine et al., 1998) have been described. To tag *HAS1* with the LacO array, 1 kb downstream of the *HAS1* ORF was PCR amplified and TOPO cloned to create pCR2.1-HAS1_3’UTR. Plasmid p6LacO128-HAS1 was made by inserting *HAS1* from pCR2.1- HAS1_3’UTR into p6LacO128 (Brickner and Walter, 2004).

### Hi-C

Cells were grown overnight, shaking at 30°C (room temperature for *S. uvarum)* in YPD media (1% yeast extract, 2% peptone, 2% dextrose), YP + raffinose (2%), or YP + galactose (2%), and for experiments with saturated culture samples, they were crosslinked at this point by resuspension and incubation in 1% formaldehyde in PBS for 20 minutes at room temperature. Crosslinking was quenched by addition of 1% solid glycine, followed by incubation for 20 minutes and a PBS wash. For arrested/synchronized experiments, fresh cultures were inoculated to OD_600_ = 0.1 in appropriate media (typically in YPD, or YP + raffinose or YP + galactose). Resultant cultures were grown at 30°C (room temperature for *S. uvarum)* until they reached OD_600_ = 0.6-0.8 at which time they were crosslinked (for exponential growth samples) or supplemented with 15μg/mL nocodazole. For arrest, samples were grown at 30°C for 2 hours following addition of drug prior to crosslinking. Arrested cultures were checked by flow cytometry. For mixture controls, samples were mixed prior to crosslinking.

Hi-C libraries were created as described (Burton et al., 2014) with the exceptions that the restriction endonuclease Sau3AI or HindIII was used to digest the chromatin and the KAPA Hyper Prep kit was used to create the Illumina library instead of the Illumina TruSeq kit. Libraries were pooled and sequenced on an Illumina NextSeq 500, with 2x80 bp reads for interspecific hybrids and 2x150 bp reads for intraspecific *S. cerevisiae* hybrids. Hi-C libraries were similar across the two restriction enzymes and biological replicates (**Figure 1—figure supplement 3**).

### Reference genomes

The *S. cerevisiae* reference annotations were downloaded from the Saccharomyces Genome Database (version R64.2.1). The *S. paradoxus* and *S. uvarum* references and annotations were downloaded from saccharomycessensustricto.org (Scannell et al., 2011) but modified to correct misassemblies evident based on synteny and Hi-C data (Figure 1—figure supplement 4). *S. paradoxus* chromosome IV was rearranged so bases 1-943,469 were followed by 1,029,253-1,193,028, then 1,027,718-1,029,252, then 943,470-1,027,717 in reverse order, followed by the remainder of the chromosome. *S. uvarum* chromosome III was rearranged so bases 219,500 onward were placed at the beginning (left end) of the chromosome, followed by the first 219,399 bases, and then new sequence determined by Sanger sequencing with primers CATTCCCATTTGTTGATTCCTG and GGATTCTATT GTT GCTAAAGGC: TAATAAGGAAGAACTGCTTATTCTTAATTATTTCTACCTACTAAACTAACTAATTATC AACAAATATCATCTATTTAATAGTATATCATCACATGCGGTGTAAGAGGATGACATA AAGATTGAGAAACAGTCATCCAGTCTAATGGAAGCTCAAATGCAAGGGCTGATAATGTAATAGGATAATGAATGACAACGTATAAAAGGAAAGAAGATAAAGCAATATTATTTTGTAGAATTATCGATTCCCTTTTGTGGATCCCTATATCCTCGAGGAGAA. S. *uvarum* chromosomes X and XII were also swapped, based on homology to *S. cerevisiae.* The *S. cerevisiae* Y12 and DBVPG6044 strain references were sequenced to 145- and 315-fold coverage using the PacBio single-molecule, real-time (SMRT) sequencing platform with P6-C2 chemistry. Each genome was assembled with FALCON (Chin et al., 2016), version June 30, 2015 hash: cee6a58, and polished with Quiver (Chin et al., 2013) version 1.1.0 to generate chromosome-length contigs (with the exclusion of chromosome XII, which was split at the rDNA array, and Y12 chromosome XIV, which was split into one large and one small contig). To call centromeres in *S. paradoxus*, we searched the region on each chromosome between the genes homologous to those nearest the centromeres in *S. cerevisiae* (e.g. *YEL001C* and *YER001W* on chromosome V) for the sequence motif N2TCAC(A/G)TGN_95-100_CCGAAN_6_ (based on an alignment of *S. uvarum, S. mikatae*, and *S. kudriavzevii* centromeres (Scannell et al., 2011)) or its reverse complement. When this motif was absent (chromosomes VII and VIII), we called the centromere as the middle 120 bp of the region. To call centromeres in Y12 and DBVPG6044, we mapped the *S. cerevisiae* S288C centromere sequences to the new references.

### Theoretical mappability analysis

Simulated reads for each hybrid genome (as in experimental data, 80 bp for interspecific hybrids and 150 bp for the intraspecific *S. cerevisiae* hybrid) were generated by taking sequences of the read length at 10 bp intervals. These reads were then remapped to the hybrid genome using bowtie2 (Langmead and Salzberg, 2012) with the —very-sensitive parameter set. The proportion of reads that mapped with mapping quality ≥ 30 to the correct location was then calculated.

### Hi-C data analysis

Sequencing reads were first pre-processed using cutadapt (Martin, 2011): reads were quality-trimmed (option -q 20), trimmed of adapter sequences, and then trimmed up to the ligation junction (if present), excluding any read pairs in which either read was shorter than 20 bp after trimming (option -m 20). The two reads in each read pair were then mapped separately using bowtie2 (Langmead and Salzberg, 2012) with the —very-sensitive parameter set. For interspecific hybrids, reads were mapped to a combined reference containing both species references, where if secondary mappings were present the best alignment must have a score ≥ 10 greater than the next best alignment. For intraspecific *S. cerevisiae* hybrids, reads were mapped separately to both strain references, keeping only read pairs mapping to both—perfectly to one reference and with ≥ 2 mutations including ≥ 1 substitution to the other. In both cases, only read pairs in which both reads mapped with MAPQ ≥ 30 were used. PCR duplicates (with identical fragment start and end positions) were removed, as were read pairs mapping within 1 kb of each other or in the same restriction fragment, which represent either unligated or invalid ligation products. The genome was then binned into 32 kb fragments (except the last fragment of each chromosome), and the number of read pairs mapping to each 32 kb genomic bin was counted based on the position of the restriction sites that were ligated together. Due to gaps in the reference genomes of *S. uvarum*, some repetitive sequences were only represented once and therefore artifactually mapped uniquely; therefore, reads mapping to annotated repetitive sequences were masked from further analysis. Similarly, gaps in the *S. paradoxus* reference led to mismapping of reads to the corresponding *S. cerevisiae* sequence; therefore, for *S. cerevisiae* x *S. paradoxus* libraries we masked regions in the *S. cerevisiae* genome where > 1 read from a *S. paradoxus* Hi-C library mapped, and vice versa. We took a similar approach to mask regions of the Y12 and DBVPG6044 references that were prone to mismapping, as estimated by haploid Y12 and DBVPG6044 Hi-C libraries. For knock-in experiments, the *HAS1pr-TDA1pr* region was masked to account for its altered genomic location. The resulting matrices were then normalized by excluding the diagonal (interactions within the same genomic bin), filtering out rows/columns with an average of less than 1 read per bin, and then multiplying each entry by the total number of read pairs divided by the column and row sums.

### Polymer model

The volume exclusion polymer model of the Rabl-like orientation was a modified version of the Tjong *et al* tethering model (Tjong et al., 2012). The beads are randomly positioned and then adjusted until all constraints are met. The model was extended from 16 chromosomes to 32, with the lengths of the *S. cerevisiae* and *S. uvarum* chromosomes. The parameters for nuclear size, centromeric constraint position and size, telomeric constraint at the nuclear periphery, and nucleolar position and size were scaled by a factor of 1.25 to reflect the roughly doubled volume of diploid nuclei (cell volume correlates with ploidy (Mortimer, 1958), and nuclear volume correlates with cell volume (Jorgensen et al., 2007)). To test the effect of smaller nuclei, all parameters were scaled by a factor of 0.8 or 0.64 from this initial diploid model. For each model, the modeling procedure was repeated 20,000 times to create a population of structures. From this population, we simulated Hi-C data by calling all beads within 45 nm of each other as contacting each other, and then counting the number of contacts between each pair of 32 kb bins. The resulting matrix of counts was normalized using the same pipeline as the experimental Hi-C data.

### Homolog proximity analysis

In order to assess homolog proximity genome-wide, we first determined which bins represented interactions between homologous sequences, and then compared the normalized interaction frequencies in those bins compared to a set of “comparable” nonhomologous bins.

In the interspecific hybrids, we determined homology by counting the number of starts or ends of one-to-one homologous gene annotations falling into each bin. Genes whose “SGD” and “BLAST” gene annotations differed were ignored. To find homologous interaction bins for genomes 1 and 2, for each bin of genome 1 we considered the bin in genome 2 where the most homologous gene ends fell to be homologous.

In the intraspecific hybrids where inter-strain mapping was much more reliable, we simulated 150 bp reads from the Y12 genome at 10 bp intervals, then mapped them to the DBVPG6044 reference. Here, for each bin of genome 1 we considered the bin in genome 2 where the most reads mapped with MAPQ ≥ 30 to be homologous.

To eliminate minor “homology” arising from repetitive sequences (e.g. telomeres), we excluded isolated homologous interaction bins lacking any other homologous interaction bins within 2 bins. To fully exclude homologous interactions from our estimates of nonhomologous interactions, any interaction bins within 2 bins of homologous interaction bins were excluded from analyses.

After determining homologous bins, we compared each homologous bin to other intergenome interactions (i.e. between chromosomes from different species/strains) involving one of the two genomic bins involved in the homologous interaction. To control for the effects of the Rabl-like orientation, we further filtered the nonhomologous interaction bins for those in which the centromeric distance (in units of 32 kb bins) was equivalent, and then for those in which the chromosome arm lengths of the two loci were within 25% of each other (in units of 32 kb bins). We also considered exclusion of the rDNA carrying chromosome XIIs as well as the centromeric bins, for which we could not fully control chromosome arm lengths. In all cases, we only considered homologous bins with at least two comparable nonhomologous bins.

To estimate genome-wide homolog proximity, we compared the sum of normalized interaction frequencies across the homologous bins to those of an equal number of randomly chosen nonhomologous bins, one comparable to each homologous bin, with replacement. We repeated this 10,000 times to obtain a distribution of genomic homolog proximity.

To obtain a view of homolog proximity strength across the genome, we compared the normalized interaction frequency in each homologous bin to the median of that in the similar nonhomologous bins, and then plotted the ratio of homologous/nonhomologous across the *S. cerevisiae* genome.

### Confocal microscopy

Cultures were grown in synthetic minimal media with 2% glucose or 2% galactose overnight at 30°C with constant shaking and harvested in log phase (OD_600_ < 0.5) or late log/stationary phase (OD_600_ x003E; 1.0). Unless noted, cultures were grown in the designated media overnight prior to imaging.

Cultures were concentrated by brief centrifugation, 1 μ l was spotted onto a microscope slide and visualized on a Leica SP5 as described (Egecioglu et al., 2014). Z-stacks of ≥ 5μm, comprising the whole yeast cell were collected. For experiments scoring peripheral localization, at least 30-50 cells were scored per biological replicate and ≥ 3 biological replicates were scored. Cells scored for peripheral localization met the following criteria 1) the strain only had one visible dot and 2) the dot was in middle third of the nucleus in the z dimension. For experiments measuring the distance between two loci we only analyzed cells in which 1) there were exactly two dots and 2) both dots were in the same z-slice or adjacent z-slices. Cells were excluded if they had abnormal nuclear morphology, only a single dot or > 2 dots.

## Microscopy data analysis

Confocal channels were merged and distances measured using LAS AF software. Distances were measured for ≥ 100 cells. Statistical tests performed include Student’s *t* test for comparisons of peripheral localization, Wilcoxon Rank Sum test for comparisons between two distributions and Fisher Exact Test for comparisons of fraction of the population < 0.55μm.

## Code availability

Code for all bioinformatic analyses is available at https://github.com/shendurelab/HybridYeastHiC.

## Data availability

GEO accession number: GSE88952

## Acknowledgements

We thank G. Yardimci, G. Bonora, M. Kircher, and K. Xue for comments and discussions, J. Andrie, J. Akey, and D. Gordon for Y12 and DBVPG6044 reference genomes and strains, and F. Winston, G. Sherlock, S. Fields, C. Payen, Y. Zheng, and D. Greig for strains. This work was supported by grants from the NIH (GM080484 to J.H.B.; P41GM103533 to M.J.D.) and NSF (1516330 to M.J.D.) J.S. is an investigator of the Howard Hughes Medical Institute. S.K. was supported by an NSF Graduate Research Fellowship. M.J.D. is a Senior Fellow in the Genetic Networks program at the Canadian Institute for Advanced Research.

## Author contributions

S.K., I.L., J.S. and M.J.D. designed the study. S.K., I.L., and D.G.B. made strains. I.L. performed Hi-C experiments. S.K. analyzed Hi-C data with assistance from K.C. D.G.B. performed microscopy. J.H.B. analyzed microscopy data. S.K. wrote the paper, with input from all authors.

## Impact Statement

Yeast homologous chromosomes exhibit condition-specific mitotic pairing at both global and local scales.

---

**Figure 1—figure supplement 1.**
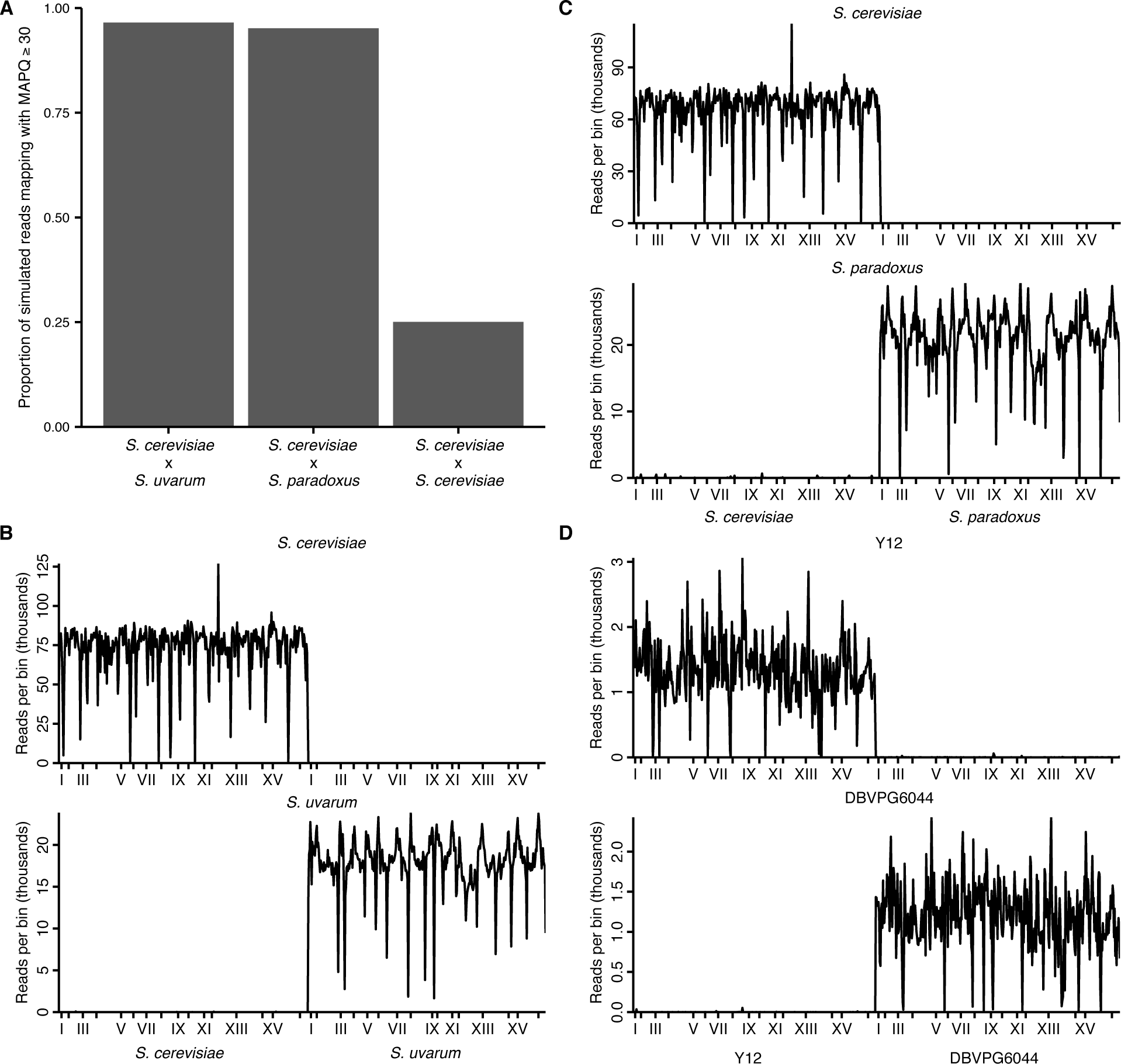
Mappability of hybrid yeast genomes. (**A**) Proportion of simulated reads from each hybrid (80 bp for interspecific and 150 bp for intraspecific hybrids, in 10 bp windows across the reference genome) that can be remapped correctly to the reference with a mapping quality score (MAPQ) of at least 30. (**B**) Number of reads (in thousands) from separate *S. cerevisiae* (left) or *S. uvarum* (right) Hi-C libraries mapping to each 32 kb genomic bin in the *S. cerevisiae* x *S. uvarum* hybrid reference genome. x ticks indicate centromeres; odd-numbered centromeres are labeled. (**C**) Same as (**B**) for *S. cerevisiae* (left) and *S. paradoxus* (right) mapping to *S. cerevisiae* x *S. paradoxus.* (**D**) Same as (**B**) for *S. cerevisiae* Y12 (left) and *S. cerevisiae* DBVPG6044 (right) mapping to Y12 x DBVPG6044.

**Figure 1—figure supplement 2.**
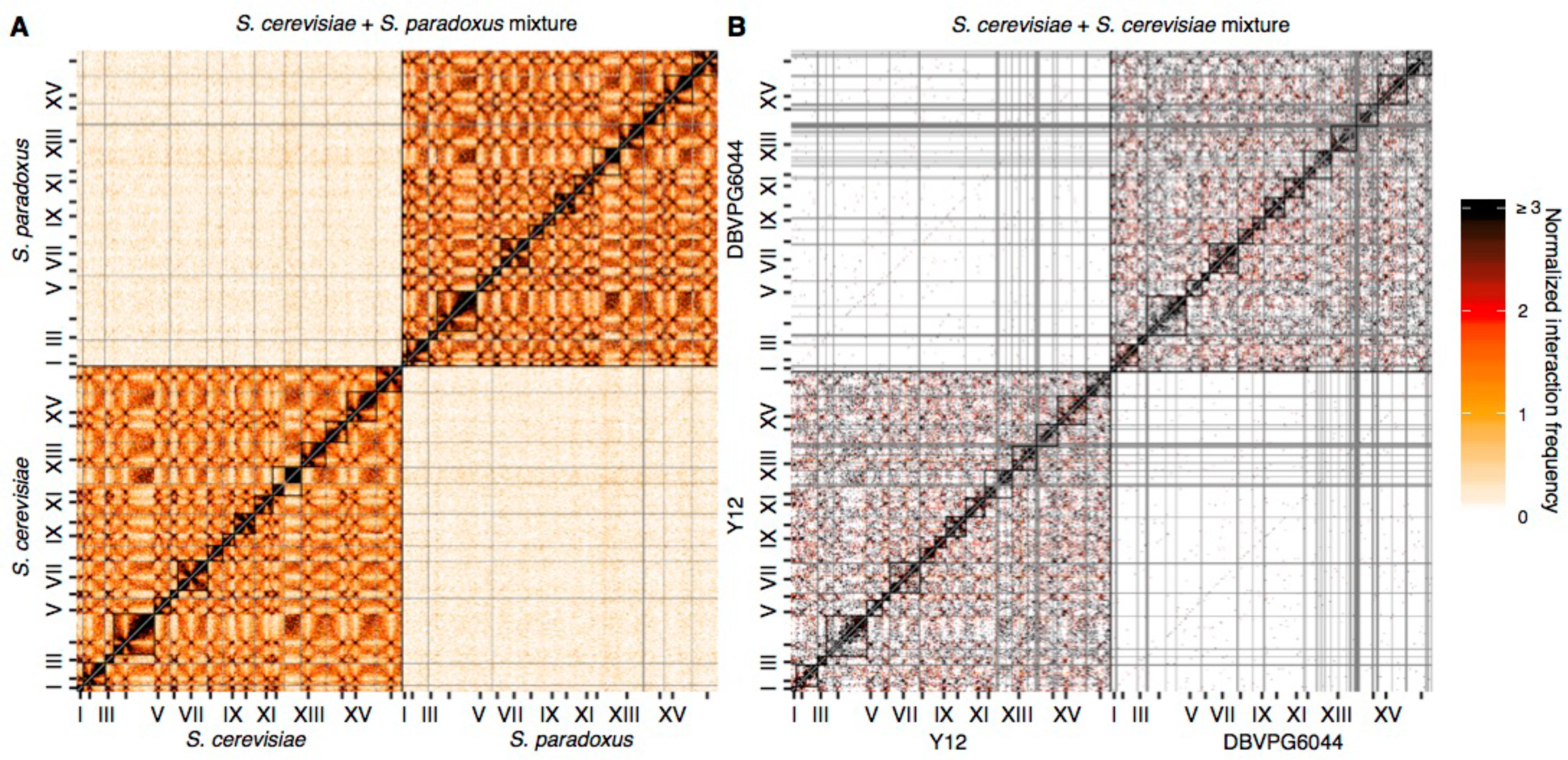
Mixture control experiments for *S. cerevisiae x S. paradoxus* and *S. cerevisiae x S. cerevisiae* hybrids. (**A**) Hi-C contact map for the *S. cerevisiae* x *S. paradoxus* hybrid in exponential growth, at 32 kb resolution. Each axis represents the *S. cerevisiae* genome followed by the *S. paradoxus* genome, in syntenic order, separated by a black line; tick marks indicate centromere positions, and odd-numbered chromosome centromeres are labeled. Intrachromosomal interactions are outlined by black squares along the diagonal. Rows and columns with an average of less than 1 read pair per bin were filtered out, and are colored grey. (**B**) Hi-C contact map for the *S. cerevisiae* Y12 x *S. cerevisiae* DBVPG6044 hybrid in exponential growth, as in (**A**).

**Figure 1—figure supplement 3.**
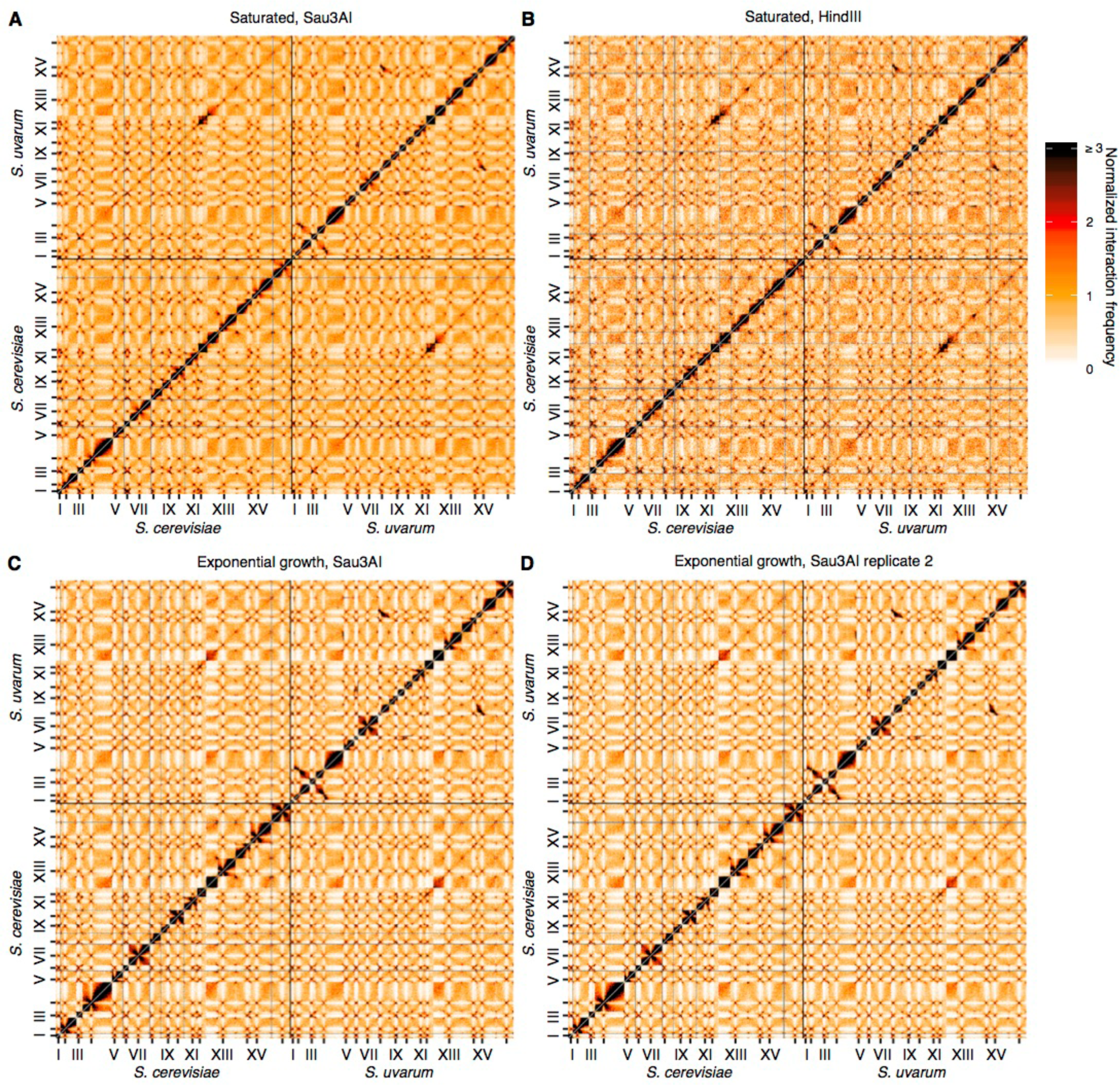
Reproducibility of Hi-C across replicates and restriction enzymes. (**A** and **B**), Hi-C contact maps at 32 kb resolution for the *S. cerevisiae* x *S. uvarum* hybrid in saturated cultures, using the restriction enzyme Sau3AI (**A**) or HindIII (**B**). Each axis represents the *S. cerevisiae* genome followed by the *S. uvarum* genome, in syntenic order, separated by a black line; ticks indicate centromeres, and odd-numbered centromeres are labeled. Rows and columns with insufficient data are colored grey. (**C and D**) Same as (**A**) and (**B**) for two biological replicates of *S. cerevisiae* x *S. uvarum* hybrid in exponential growth, using the restriction enzyme Sau3AI.

**Figure 1—figure supplement 4.**
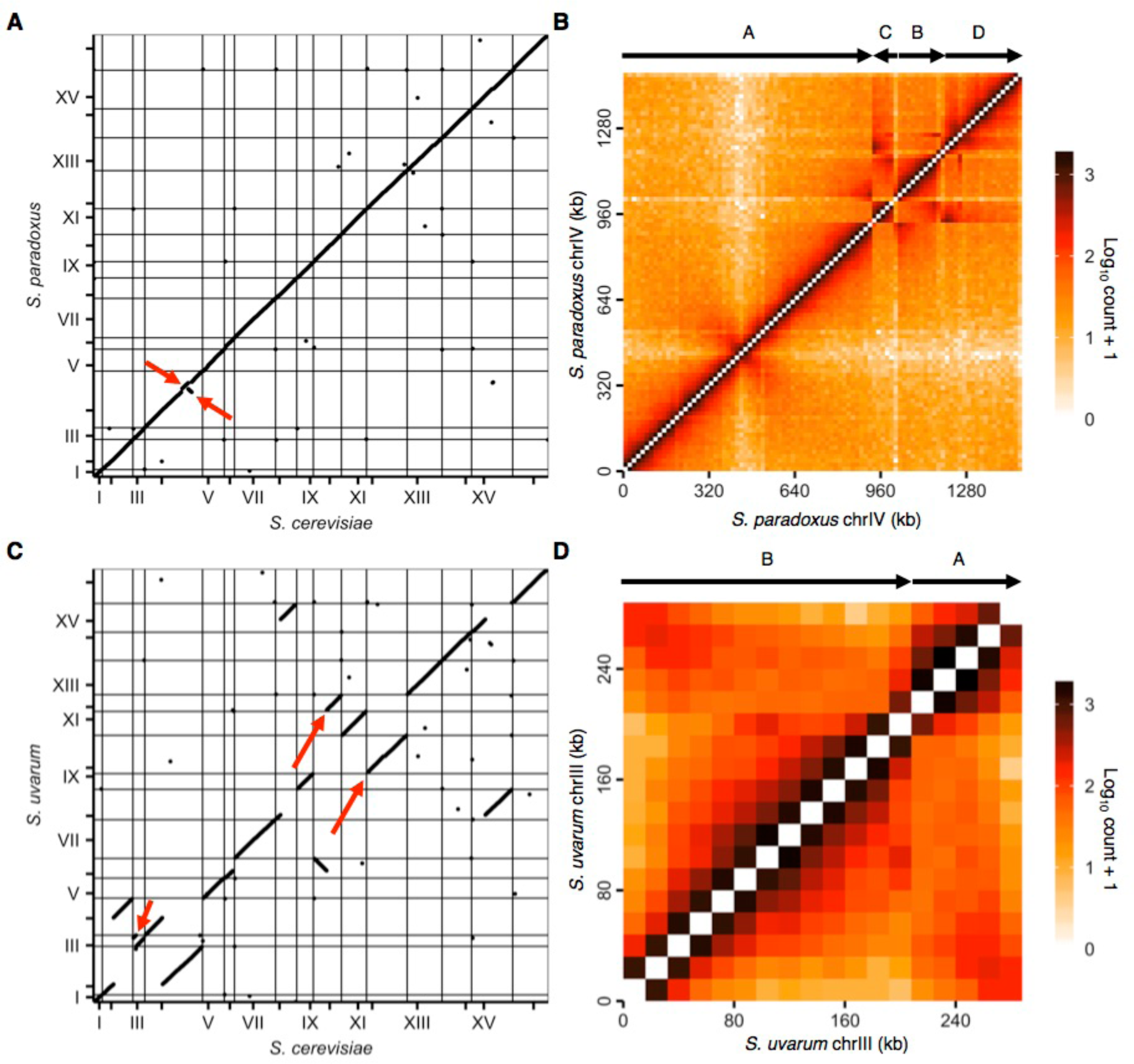
Revisions to *S. paradoxus* and *S. uvarum* reference genomes. (**A**) Start and end positions of homologous genes on the original *S. cerevisiae* and *S. paradoxus* reference genomes. Each point represents the start or end of a homologous gene pair. Horizontal and vertical lines indicate starts of chromosomes, and ticks indicate centromere positions. Odd numbered chromosomes are numbered. Red arrows indicate unexpected rearrangements. (**B**) Raw contact map for *S. paradoxus* chromosome IV using the original reference genome. Arrows above the heat map represent segments of the chromosome, labeled A-D in order and orientation of synteny. (**C**) Same as (**A**) for *S. cerevisiae* and *S. uvarum.* Red arrows indicate an unexpected rearrangement in chromosome III, and *S. uvarum* chromosomes X and XII, which are homologous to *S. cerevisiae* chromosome XII and X, respectively. (**D**) Same as (**B**) for *S. uvarum* chromosome III.

**Figure 2—figure supplement 1.**
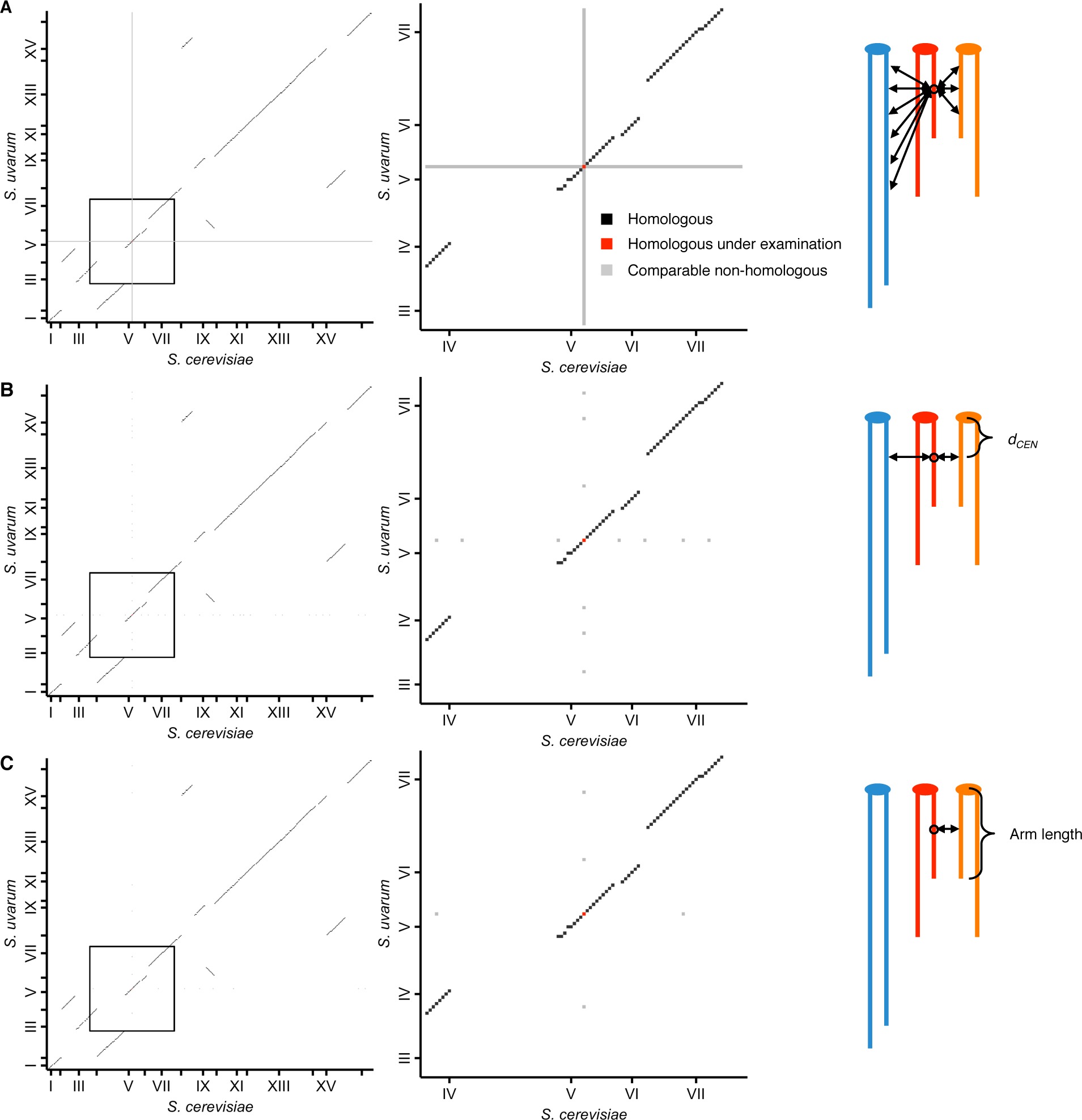
Schematic of homolog proximity analysis. Representation of how homologous interactions (black squares) were compared to various subsets of nonhomologous interactions (grey squares), either including all interactions with either homologous locus (red squares) (A), restricted to interactions with loci at a similar centromeric distance, or *d*_*CEN*_ (same number of 32 kb bins) (B), or restricted to interactions with loci at a similar centromeric distance and on a chromosome arm of similar length (within 25%) (C). Left panels show all interactions between the *S. cerevisiae* and *S. uvarum* genomes; middle panels show enlarged view of the area outlined in the left panels. Tick marks indicate centromere positions. Right panels represent the nonhomologous interactions being used for comparison; different colors represent nonhomologous chromosomes, and double-headed arrows represent interactions with the locus of interest (circled).

**Figure 2—figure supplement 2.**
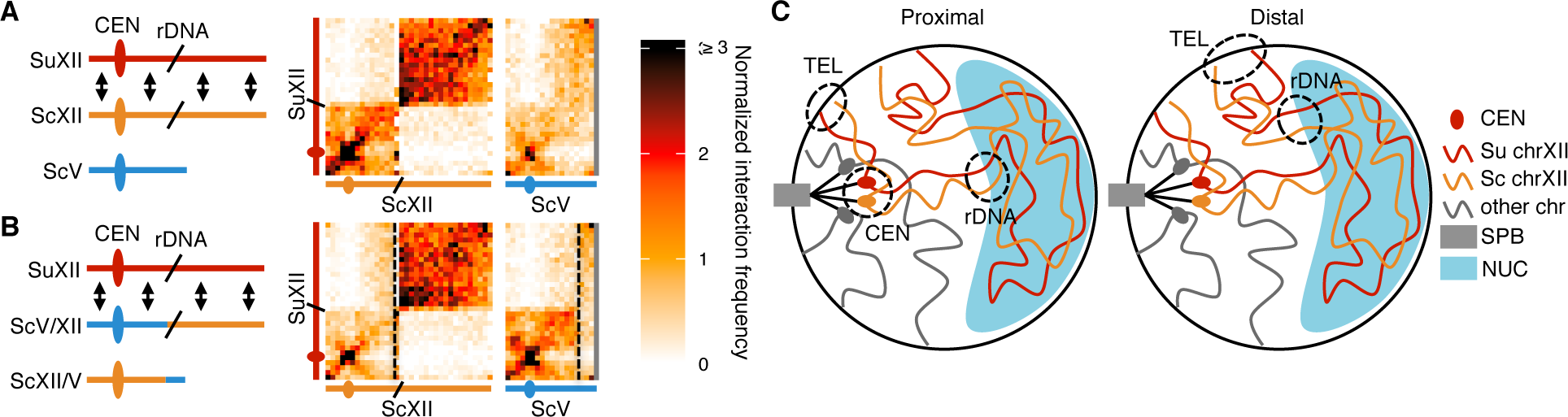
rDNA-carrying chromosomes interact preferentially due to shared tethering. (**A** and **B**) Schematics and contact maps of rDNA-carrying chromosomes *S. uvarum* chromosome XII (SuXII) and *S. cerevisiae* chromosome XII (ScXII), and *S. cerevisiae* chromosome V (ScV), in wild-type *S. cerevisiae* x *S. uvarum* hybrids (**A**) and a strain with a translocation between ScXII and ScV (**B**), both in exponential growth. Ovals indicate centromeres and slanted lines indicate the rDNA arrays. Double-headed arrows indicate enhanced interactions. Dashed lines in the contact maps indicate the translocation breakpoints. (**C**) Schematic of how the rDNA-carrying chromosomes *S. cerevisiae* (Sc) chrXII and *S. uvarum* (Su) chrXII preferentially interact due to shared tethering. The proximal halves (left diagram) of the chromosomes, which contain the centromeres (CEN), are tethered at the spindle pole body (SPB) at their centromeres, at the periphery at their telomeres (TEL), and at the nucleolus (NUC) at their rDNA arrays (rDNA). The distal halves (right diagram) are tethered at their telomeres and rDNA, but not their centromeres. These combinations of tethering points are not found in other chromosomes (shown in grey). Su, *S. uvarunm;* Sc, *S. cerevisiae.*

**Figure 3—figure supplement 1.**
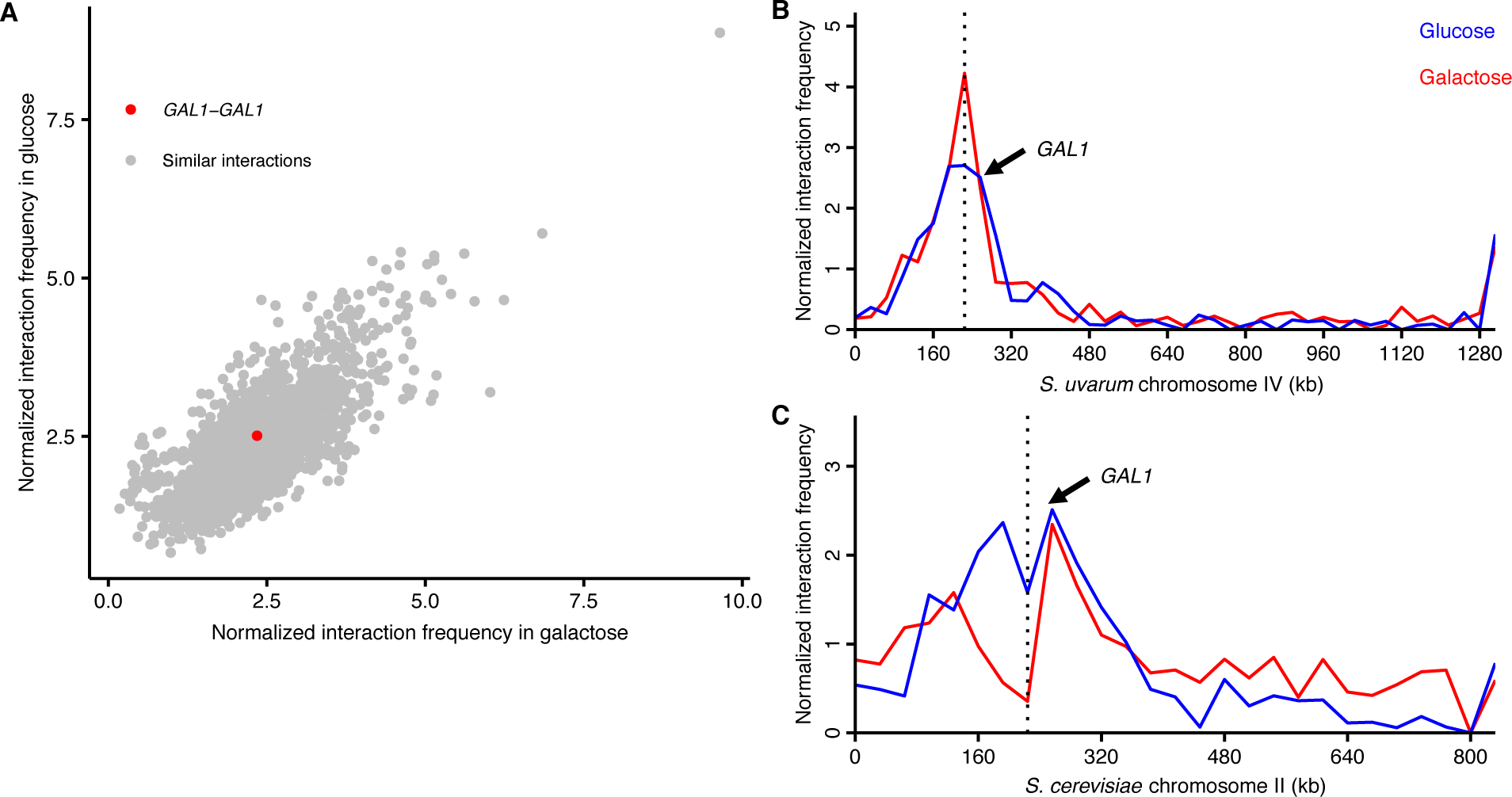
*GAL1* homologs do not detectably pair during galactose induction. (**A**) Scatter plot of normalized interaction frequencies in galactose (x-axis) and in glucose *(y-*axis), between the *GAL1* homologs (in red) and other interactions between loci at the same centromeric distance (in grey). All interactions are between 32 kb bins. (**B**) Normalized interaction frequencies between the *S. cerevisiae GAL1* and the *S. uvarum* chromosome IV in glucose (blue) and galactose (red). Arrow points toward interaction between *GAL1* homologs. (**C**) Same as (**B**) for the *S. uvarum GAL1* and the *S. cerevisiae* chromosome II.

**Figure 4—figure supplement 1.**
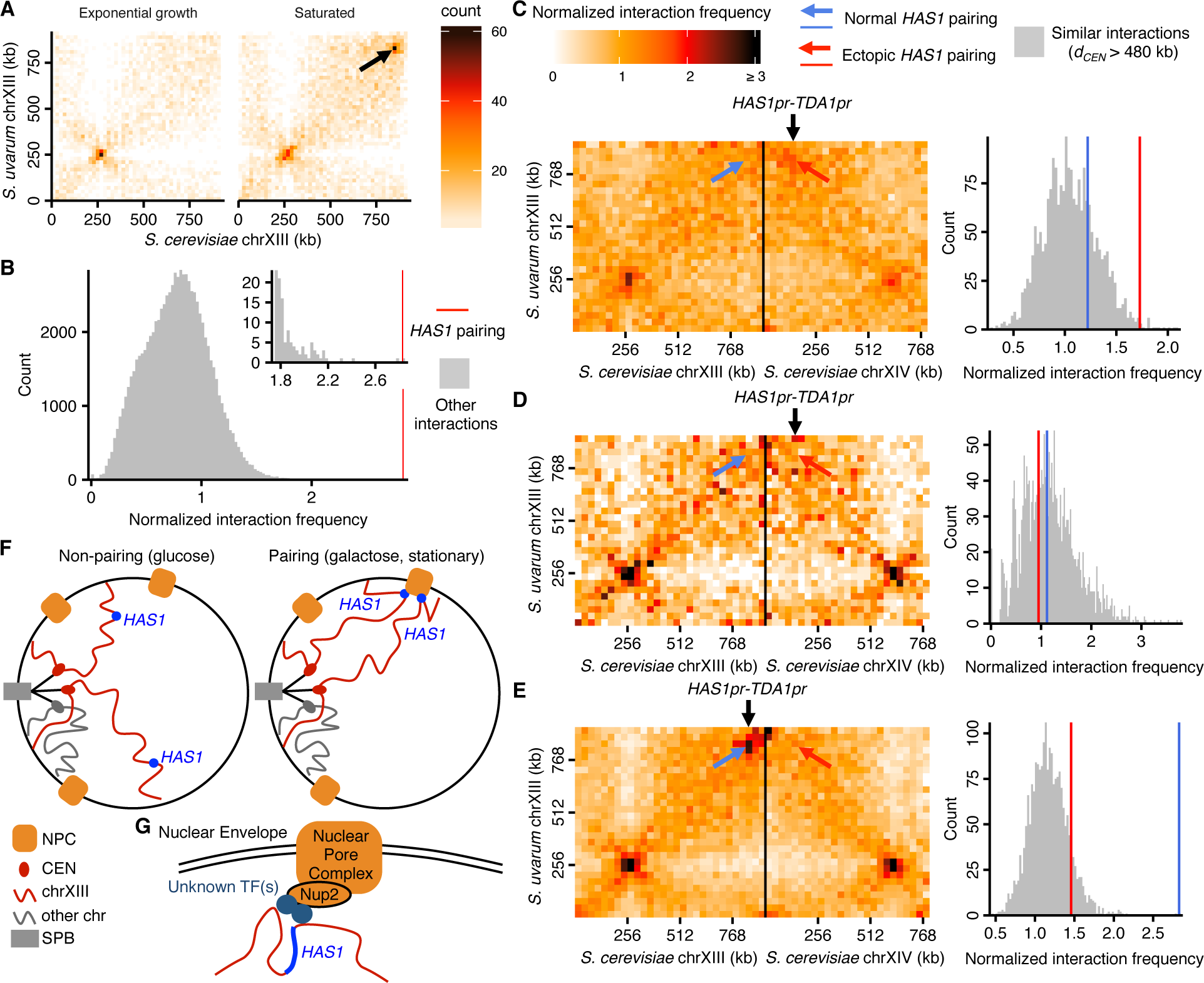
Exceptional inducible homolog pairing at *HAS1* locus is recapitulated ectopically by the *HAS1* promoter and mediated by nuclear pores. (**A**) Raw Hi-C contact maps at 20 kb resolution for *S. cerevisiae* and *S. uvarum* chromosome XIII. Arrow points to strongest interaction in saturated culture excluding regions near centromeres, at positions 840,000–860,000 on the *S. cerevisiae* chromosome XIII, the target of deletion studies. (**B**) Histogram comparing *HAS1* homologous pairing interaction frequency (red line) to all other interactions (grey) between an *S. cerevisiae* locus and an *S. uvarum* locus (32 kb bin), with both loci ≥ 3 bins from a centromere, > 1 bin from a telomere, and not both on an rDNA-carrying chromosome. Inset shows enlarged view of right end of plot. (**C, D**, and **E**) Genomic relocation of the *S. cerevisiae HAS1pr-TDA1pr* 1 kb region causes inducible ectopic pairing with the *S. uvarum HAS1* allele. (Left) Hi-C contact maps of interactions between *S. uvarum* chromosome XIII and *S. cerevisiae* chromosomes XIII and XIV in the *S. cerevisiae* x *S. uvarum* hybrid with the *S. cerevisiae HAS1pr-TDA1pr* region moved to *S. cerevisiae* chromosome XIV (location indicated by vertical black arrow), in saturated culture (**C**) and in exponential growth (**D**), compared to the wild-type hybrid (E) in saturated culture. Interactions between the *S. uvarum HAS1* locus and the original *S. cerevisiae HAS1* locus are indicated by a blue arrow, whereas interactions with the new *HAS1pr-TDA1pr* locus are indicated by a red arrow. (Right) Histograms comparing the pairing frequency of *S. uvarum HAS1* and either the normal *S. cerevisiae HAS1* locus (blue line) or the ectopic *HAS1* locus (red line) to the frequency of all similar interactions, i.e. those between an *S. cerevisiae* locus and an *S. uvarum* locus (32 kb bin), with both loci ≥ 15 bins from a centromere (or *d*_*CEN*_ > 480 kb), > 1 bin from a telomere, and not both on an rDNA-carrying chromosome. (**F**) Schematic of how nuclear pore association mediates homologous *HAS1* pairing. NPC, nuclear pore complex; CEN, centromere; chr, chromosome; SPB, spindle pole body. (**G**) Model of nuclear pore association with *HAS1* locus. TF, transcription factor.

## Supplementary Files

**Supplementary Table 1.**
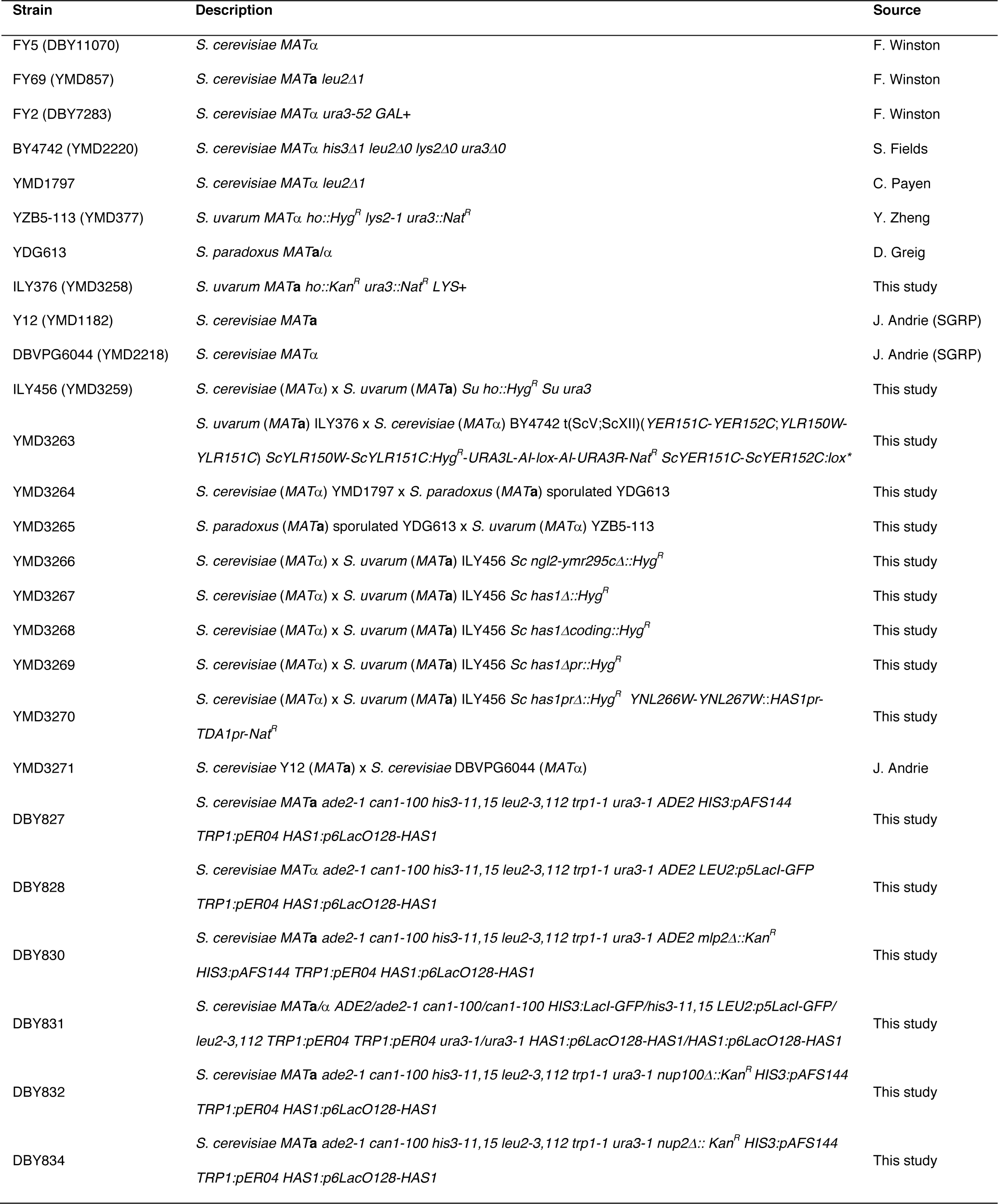
Strains used in this study.

| Strain | Description | Source |
| --- | --- | --- |
| FY5 (DBY11070) | *S. cerevisiae* MATα | F. Winston |
| FY69 (YMD857) | *S. cerevisiae* MATa *leu2Δ1* | F. Winston |
| FY2 (DBY7283) | *S. cerevisiae* MATα *ura3-52 GAL+* | F. Winston |
| BY4742 (YMD2220) | *S. cerevisiae* MATα *his3Δ1 leu2Δ0 lys2Δ0 ura3Δ0* | S. Fields |
| YMD1797 | *S. cerevisiae* MATα *leu2Δ1* | C. Payen |
| YZB5-113 (YMD377) | *S. uvarum* MATα *ho::Hyg<sup>R</sup> lys2-1 ura3::Nat<sup>R</sup>* | Y. Zheng |
| YDG613 | *S. paradoxus* MATa/$\alpha$ | D. Greig |
| ILY376 (YMD3258) | *S. uvarum* MATa *ho::Kan<sup>R</sup> ura3::Nat<sup>R</sup> LYS+* | This study |
| Y12 (YMD1182) | *S. cerevisiae* MATa | J. Andrie (SGRP) |
| DBVPG6044 (YMD2218) | *S. cerevisiae* MATα | J. Andrie (SGRP) |
| ILY456 (YMD3259) | *S. cerevisiae* (MATα) x *S. uvarum* (MATa) *Su ho::Hyg<sup>R</sup> Su ura3* | This study |
| YMD3263 | *S. uvarum* (MATa) ILY376 x *S. cerevisiae* (MATα) BY4742 t(ScV;ScXII)(*YER151C-YER152C;YLR150W-YLR151C*) *ScYLR150W-ScYLR151C:Hyg<sup>R</sup>-URA3L-AI-lox-AI-URA3R-Nat<sup>R</sup> ScYER151C-ScYER152C:lox\** | This study |
| YMD3264 | *S. cerevisiae* (MATα) YMD1797 x *S. paradoxus* (MATa) sporulated YDG613 | This study |
| YMD3265 | *S. paradoxus* (MATa) sporulated YDG613 x *S. uvarum* (MATα) YZB5-113 | This study |
| YMD3266 | *S. cerevisiae* (MATα) x *S. uvarum* (MATa) ILY456 *Sc ngl2-ymr295cΔ::Hyg<sup>R</sup>* | This study |
| YMD3267 | *S. cerevisiae* (MATα) x *S. uvarum* (MATa) ILY456 *Sc has1Δ::Hyg<sup>R</sup>* | This study |
| YMD3268 | *S. cerevisiae* (MATα) x *S. uvarum* (MATa) ILY456 *Sc has1Δcoding::Hyg<sup>R</sup>* | This study |
| YMD3269 | *S. cerevisiae* (MATα) x *S. uvarum* (MATa) ILY456 *Sc has1Δpr::Hyg<sup>R</sup>* | This study |
| YMD3270 | *S. cerevisiae* (MATα) x *S. uvarum* (MATa) ILY456 *Sc has1prΔ::Hyg<sup>R</sup> YNL266W-YNL267W::HAS1pr-TDA1pr-Nat<sup>R</sup>* | This study |
| YMD3271 | *S. cerevisiae* Y12 (MATa) x *S. cerevisiae* DBVPG6044 (MATα) | J. Andrie |
| DBY827 | *S. cerevisiae* MATa *ade2-1 can1-100 his3-11,15 leu2-3,112 trp1-1 ura3-1 ADE2 HIS3:pAFS144 TRP1:pER04 HAS1:p6LacO128-HAS1* | This study |
| DBY828 | *S. cerevisiae* MATα *ade2-1 can1-100 his3-11,15 leu2-3,112 trp1-1 ura3-1 ADE2 LEU2:p5LacI-GFP TRP1:pER04 HAS1:p6LacO128-HAS1* | This study |
| DBY830 | *S. cerevisiae* MATa *ade2-1 can1-100 his3-11,15 leu2-3,112 trp1-1 ura3-1 ADE2 mlp2Δ::Kan<sup>R</sup> HIS3:pAFS144 TRP1:pER04 HAS1:p6LacO128-HAS1* | This study |
| DBY831 | *S. cerevisiae* MATa/$\alpha$ *ADE2/ade2-1 can1-100/can1-100 HIS3:LacI-GFP/his3-11,15 LEU2:p5LacI-GFP/leu2-3,112 TRP1:pER04 TRP1:pER04 ura3-1/ura3-1 HAS1:p6LacO128-HAS1/HAS1:p6LacO128-HAS1* | This study |
| DBY832 | *S. cerevisiae* MATa *ade2-1 can1-100 his3-11,15 leu2-3,112 trp1-1 ura3-1 nup100Δ::Kan<sup>R</sup> HIS3:pAFS144 TRP1:pER04 HAS1:p6LacO128-HAS1* | This study |
| DBY834 | *S. cerevisiae* MATa *ade2-1 can1-100 his3-11,15 leu2-3,112 trp1-1 ura3-1 nup2Δ::Kan<sup>R</sup> HIS3:pAFS144 TRP1:pER04 HAS1:p6LacO128-HAS1* | This study |

SGRP, *Saccharomyces* Genome Resequencing Project (Liti et al., 2009)

SGRP, *Saccharomyces* Genome Resequencing Project (Liti et al., 2009)

**Supplementary Table 2.** Hi-C libraries.

| Strain(s) | Condition | Restriction Enzyme | Read Length | Read pairs |
| --- | --- | --- | --- | --- |
| YMD1797 | raffinose | Sau3AI | 80 bp | 50,609,269 |
| YDG613 | exponential | Sau3AI | 80 bp | 17,996,489 |
| YZB5-113 | exponential | Sau3AI | 80 bp | 20,160,361 |
| Y12 | exponential | Sau3AI | 150 bp | 70,141,448 |
| DBVPG6044 | exponential | Sau3AI | 150 bp | 62,960,123 |
| FY69 + YZB5-113 | saturated | Sau3AI | 80 bp | 17,238,325 |
| FY69 + YDG613 | exponential | Sau3AI | 80 bp | 16,280,645 |
| Y12 + DBVPG6044 | exponential | Sau3AI | 150 bp | 63,418,065 |
| ILY456 | saturated | Sau3AI | 80 bp | 25,139,616 |
| ILY456 | saturated | HindIII | 80 bp | 8,268,022 |
| ILY456 | exponential | Sau3AI | 80 bp | 12,131,350 |
| ILY456 | exponential, rep 2 | Sau3AI | 80 bp | 20,007,389 |
| ILY456 | galactose | Sau3AI | 80 bp | 18,384,435 |
| ILY456 | nocodazole | Sau3AI | 80 bp | 24,915,498 |
| YMD3263 | saturated | Sau3AI | 80 bp | 25,866,783 |
| YMD3263 | exponential | Sau3AI | 80 bp | 26,592,214 |
| YMD3264 | saturated | Sau3AI | 80 bp | 62,902,987 |
| YMD3264 | exponential | Sau3AI | 80 bp | 24,502,214 |
| YMD3265 | saturated | Sau3AI | 80 bp | 41,615,076 |
| YMD3265 | exponential | Sau3AI | 80 bp | 28,280,637 |
| YMD3266 | saturated | Sau3AI | 80 bp | 23,981,947 |
| YMD3267 | saturated | Sau3AI | 80 bp | 23,045,668 |
| YMD3268 | saturated | Sau3AI | 80 bp | 13,268,816 |
| YMD3269 | saturated | Sau3AI | 80 bp | 11,219,451 |
| YMD3270 | saturated | Sau3AI | 80 bp | 18,929,190 |
| YMD3270 | exponential | Sau3AI | 80 bp | 14,620,886 |
| YMD3271 | saturated | Sau3AI | 150 bp | 73,586,493 |
| YMD3271 | exponential | Sau3AI | 150 bp | 64,048,246 |
| TOTAL |  |  |  | 880,111,643 |

## References

Ahmed, S., Brickner, D.G., Light, W.H., Cajigas, I., McDonough, M., Froyshteter, A.B., Volpe, T., and Brickner, J.H. (2010). DNA zip codes control an ancient mechanism for gene targeting to the nuclear periphery. Nat. Cell Biol. 12, 111–118.

Brickner, J.H., and Walter, P. (2004). Gene Recruitment of the Activated INO1 Locus to the Nuclear Membrane. PLoS Biol. 2, e342.

Brickner, D.G., Ahmed, S., Meldi, L., Thompson, A., Light, W., Young, M., Hickman, T.L., Chu, F., Fabre, E., and Brickner, J.H. (2012). Transcription factor binding to a DNA zip code controls interchromosomal clustering at the nuclear periphery. Dev. Cell 22, 1234–1246.

Brickner, D.G., Sood, V., Tutucci, E., Coukos, R., Viets, K., Singer, R., and Brickner, J.H. (2016). Subnuclear positioning and interchromosomal clustering of the GAL1–10 locus are controlled by separable, interdependent mechanisms. Mol. Biol. Cell mbc.E16-03-0174.

Burgess, S.M., and Kleckner, N. (1999). Collisions between yeast chromosomal loci in vivo are governed by three layers of organization. Genes Dev. 13, 1871–1883.

Burgess, S.M., Kleckner, N., and Weiner, B.M. (1999). Somatic pairing of homologs in budding yeast: existence and modulation. Genes Dev. 13, 1627–1641.

Burton, J.N., Liachko, I., Dunham, M.J., and Shendure, J. (2014). Species-level deconvolution of metagenome assemblies with Hi-C-based contact probability maps. G3 (Bethesda). 4, 1339–1346.

Casolari, J.M., Brown, C.R., Komili, S., West, J., Hieronymus, H., and Silver, P.A. (2004). Genome-Wide Localization of the Nuclear Transport Machinery Couples Transcriptional Status and Nuclear Organization. Cell 117, 427–439.

Chin, C.-S., Alexander, D.H., Marks, P., Klammer, A.A., Drake, J., Heiner, C., Clum, A., Copeland, A., Huddleston, J., Eichler, E.E., et al. (2013). Nonhybrid, finished microbial genome assemblies from long-read SMRT sequencing data. Nat. Methods 10, 563–569.

Chin, C.-S., Peluso, P., Sedlazeck, F.J., Nattestad, M., Concepcion, G.T., Clum, A., Dunn, C., O’Malley, R., Figueroa-Balderas, R., Morales-Cruz, A., et al. (2016). Phased diploid genome assembly with single-molecule real-time sequencing. Nat. Methods.

Dekker, J., Rippe, K., Dekker, M., and Kleckner, N. (2002). Capturing chromosome conformation. Science 295, 1306–1311.

Dieppois, G., Iglesias, N., and Stutz, F. (2006). Cotranscriptional recruitment to the mRNA export receptor Mex67p contributes to nuclear pore anchoring of activated genes. Mol. Cell. Biol. 26, 7858–7870.

Duan, Z., Andronescu, M., Schutz, K., McIlwain, S., Kim, Y.J., Lee, C., Shendure, J., Fields, S., Blau, C.A., and Noble, W.S. (2010). A three-dimensional model of the yeast genome. Nature 465, 363–367.

Dultz, E., Tjong, H., Weider, E., Herzog, M., Young, B., Brune, C., Müllner, D., Loewen, C., Alber, F., and Weis, K. (2016). Global reorganization of budding yeast chromosome conformation in different physiological conditions. J. Cell Biol. 212, 321–334.

Egecioglu, D.E., D’Urso, A., Brickner, D.G., Light, W.H., and Brickner, J.H. (2014). Approaches to studying subnuclear organization and gene-nuclear pore interactions. Methods Cell Biol. 122, 463–485.

Fischer, G., James, S. a, Roberts, I.N., Oliver, S.G., and Louis, E.J. (2000). Chromosomal evolution in Saccharomyces. Nature 405, 451–454.

González, S.S., Barrio, E., Gafner, J., and Querol, A. (2006). Natural hybrids from Saccharomyces cerevisiae, Saccharomyces bayanus and Saccharomyces kudriavzevii in wine fermentations. FEMS Yeast Res. 6, 1221–1234.

Guidi, M., Ruault, M., Marbouty, M., Loïodice, I., Cournac, A., Billaudeau, C., Hocher, A., Mozziconacci, J., Koszul, R., and Taddei, A. (2015). Spatial reorganization of telomeres in long-lived quiescent cells. Genome Biol. 16, 206.

Güldener, U., Heck, S., Fielder, T., Beinhauer, J., and Hegemann, J.H. (1996). A new efficient gene disruption cassette for repeated use in budding yeast. Nucleic Acids Res. 24, 2519–2524.

Jin, Q., Trelles-Sticken, E., Scherthan, H., and Loidl, J. (1998). Yeast nuclei display prominent centromere clustering that is reduced in nondividing cells and in meiotic prophase. J. Cell Biol. 141, 21–29.

Jorgensen, P., Edgington, N.P., Schneider, B.L., Rupes, I., Tyers, M., and Futcher, B. (2007). The size of the nucleus increases as yeast cells grow. Mol. Biol. Cell 18, 3523–3532.

Kellis, M., Patterson, N., Endrizzi, M., Birren, B., and Lander, E.S. (2003). Sequencing and comparison of yeast species to identify genes and regulatory elements. Nature 423, 241–254.

Langmead, B., and Salzberg, S.L. (2012). Fast gapped-read alignment with Bowtie 2. Nat. Methods 9, 357–359.

Levy, S.F., Blundell, J.R., Venkataram, S., Petrov, D.A., Fisher, D.S., and Sherlock, G. (2015). Quantitative evolutionary dynamics using high-resolution lineage tracking. Nature 519, 181–186.

Liti, G., Carter, D.M., Moses, A.M., Warringer, J., Parts, L., James, S.A., Davey, R.P., Roberts, I.N., Burt, A., Koufopanou, V., et al. (2009). Population genomics of domestic and wild yeasts. Nature 458, 337–341.

Longtine, M.S., Mckenzie III, A., Demarini, D.J., Shah, N.G., Wach, A., Brachat, A., Philippsen, P., and Pringle, J.R. (1998). Additional modules for versatile and economical PCR-based gene deletion and modification in Saccharomyces cerevisiae. Yeast 14, 953–961.

Lorenz, A., Fuchs, J., Bürger, R., and Loidl, J. (2003). Chromosome pairing does not contribute to nuclear architecture in vegetative yeast cells. Eukaryot. Cell 2, 856–866.

Luthra, R., Kerr, S.C., Harreman, M.T., Apponi, L.H., Fasken, M.B., Ramineni, S., Chaurasia, S., Valentini, S.R., and Corbett, A.H. (2007). Actively transcribed GAL genes can be physically linked to the nuclear pore by the SAGA chromatin modifying complex. J. Biol. Chem. 282, 3042–3049.

Martin, M. (2011). Cutadapt removes adapter sequences from high-throughput sequencing reads. Embnet.journal 17, 10–12.

Mertens, S., Steensels, J., Saels, V., De Rouck, G., Aerts, G., and Verstrepen, K.J. (2015). A large set of newly created interspecific Saccharomyces hybrids increases aromatic diversity in lager beers. Appl. Environ. Microbiol. 81, 8202–8214.

Metz, C.W. (1916). Chromosome studies on the Diptera. II. The paired association of chromosomes in the Diptera, and its significance. J. Exp. Zool. 21, 213–279.

Mortimer, R.K. (1958). Radiobiological and Genetic Studies on a Polyploid Series (Haploid to Hexaploid) of Saccharomyces cerevisiae. Radiat. Res. 9, 312.

Randise-Hinchliff, C., Coukos, R., Sood, V., Sumner, M.C., Zdraljevic, S., Meldi Sholl, L., Garvey Brickner, D., Ahmed, S., Watchmaker, L., and Brickner, J.H. (2016). Strategies to regulate transcription factor-mediated gene positioning and interchromosomal clustering at the nuclear periphery. J. Cell Biol. 212, 633–646.

Rutledge, M.T., Russo, M., Belton, J.-M., Dekker, J., and Broach, J.R. (2015). The yeast genome undergoes significant topological reorganization in quiescence. Nucleic Acids Res. 43, 8299–8313.

Scannell, D.R., Zill, O.A., Rokas, A., Payen, C., Dunham, M.J., Eisen, M.B., Rine, J., Johnston, M., and Hittinger, C.T. (2011). The Awesome Power of Yeast Evolutionary Genetics: New Genome Sequences and Strain Resources for the Saccharomyces sensu stricto Genus. G3 1, 11–25.

Straight, A.F., Belmont, A.S., Robinett, C.C., and Murray, A.W. (1996). GFP tagging of budding yeast chromosomes reveals that protein-protein interactions can mediate sister chromatid cohesion. Curr. Biol. 6, 1599–1608.

Taddei, A., Van Houwe, G., Hediger, F., Kalck, V., Cubizolles, F., Schober, H., and Gasser, S.M. (2006). Nuclear pore association confers optimal expression levels for an inducible yeast gene. Nature 441, 774–778.

Taddei, A., Schober, H., and Gasser, S.M. (2010). The Budding Yeast Nucleus. Cold Spring Harb. Perspect. Biol. 2, a000612.

Therizols, P., Fairhead, C., Cabal, G.G., Genovesio, A., Olivo-Marin, J.-C., Dujon, B., and Fabre, E. (2006). Telomere tethering at the nuclear periphery is essential for efficient DNA double strand break repair in subtelomeric region. J. Cell Biol. 172, 189–199.

Tjong, H., Gong, K., Chen, L., and Alber, F. (2012). Physical tethering and volume exclusion determine higher-order genome organization in budding yeast. Genome Res. 22, 1295–1305.

Wong, H., Marie-Nelly, H., Herbert, S., Carrivain, P., Blanc, H., Koszul, R., Fabre, E., and Zimmer, C. (2012). A Predictive Computational Model of the Dynamic 3D Interphase Yeast Nucleus. Curr. Biol. 22, 1881–1890.

Yang, C.H., Lambie, E.J., Hardin, J., Craft, J., and Snyder, M. (1989). Higher order structure is present in the yeast nucleus: autoantibody probes demonstrate that the nucleolus lies opposite the spindle pole body. Chromosoma 98, 123–128.

